# Structural analysis of DNA binding and flexibility of the homohexameric AAA+ helicase SaPIbov1 Rep

**DOI:** 10.64898/2026.11.19.767935

**Authors:** Heewhan Shin, Ignacio Mir-Sanchis, Ying Pigli, Phoebe A. Rice

## Abstract

The replication initiation proteins (Reps) encoded by *Staphylococcus aureus* pathogenicity islands (SaPIs) are helicases that recognize their own origins of replication (oris). Despite their ability to melt the cognate ori without accessory proteins, the nature of the transition from origin-loaded to actively unwinding states remains poorly understood. Here, we report crystal and cryo-EM structures of the SaPIbov1 Rep, a homohexameric superfamily 6 (SF6) helicase, in apo and nucleotide plus duplex DNA-bound states. Comparison to superfamily 3 (SF3) SaPI Reps and archaeal/eukaryotic MCMs reveals differing structural adaptations for DNA engagement. Notably, SaPIbov1 Rep retains pore-loop 1 (PL1) and the pre-sensor 1 β-hairpin (PS1) for DNA-binding, but lacks the Helix-2 insertion (H2i) conserved in MCM helicases and instead possesses an elongated PS1 providing extended DNA contact. 3D variability analysis (3DVA) resolved multiple ring-open conformational intermediate states, with ring asymmetry correlating to DNA binding. The duplex-bound single hexamer captured here likely represents a pre-replicative intermediate preceding double hexamer assembly. We also show that the disordered N-terminus of SaPIbov1 Rep binds its cognate primase and predict that the primase recruits the host DNA polymerase III.

## Introduction

Hexameric helicases of the AAA+ (ATPase Associated with various cellular Activities) superfamily couple ATP hydrolysis to DNA unwinding, providing single-stranded templates essential for genome maintenance across all domains of life and in viruses (1). Despite evolutionary divergence, AAA+ proteins share a conserved architecture in which large and small AAA+ domains assemble composite active sites at subunit interfaces (2–4). The large AAA+ domain contributes the conserved catalytic residues of the Walker A/B motifs that bind the nucleotide cofactor in *cis*, and an arginine finger from an adjacent subunit that contacts the ɣ-phosphate in *trans* (5, 6). The sensor II residue, located in the small AAA+ lid domain further contributes to ɣ-phosphate coordination, acting either in *cis*- or in *trans*-, depending on its structural arrangement (4, 7).

Most replicative helicases assemble into nucleotide-bound, ring-shaped hexamers typically arranged in a right-handed spiral staircase architecture, with a central channel through which DNA is threaded (8, 9). Four to six subunits simultaneously engage DNA through conserved DNA binding motifs, a configuration shared across bacterial DnaB (RecA-like) (10, 11), viral SF3 helicases (papillomavirus E1, SV40 LTag) (8, 12, 13), and archaeal and eukaryotic MCMs (14–20). Despite many similarities, bacterial replicative helicases generally translocate along the lagging strand in the 5’ to 3’ direction, while eukaryotic and archaeal MCM helicases translocate along the leading strand in the 3’ to 5’ direction (21). In both groups, nucleotide occupancy varies around the ring: ATP (or a non-hydrolyzable ATP analog) occupies the higher staircase positions, while ADP-bound or apo states predominate at lower positions (9). This gradient correlates with variations in intersubunit geometry, where ATP-bound interfaces are tighter and ADP- or apo interfaces are looser (12, 20, 22–24). A similar spiral staircase architecture is also seen in AAA+ protein unfoldases such as ClpX (25–29), which thread extended protein chains instead of DNA. Though exceptions exist, spiral architectures generally reflect active translocation intermediates, while planar rings represent pre- or post-translocation states (8, 9, 11).

Two broad translocation models have been proposed for hexameric helicases. In the hand-over-hand mechanism, supported by biochemical and structural studies for bacterial and viral hexameric helicases including RecA-family, bacteriophage T7 gp4, and T4 gp41 (8, 9, 11), each subunit cycles through ATP binding, hydrolysis, and ADP release in a coordinated sequential fashion: the subunit at the bottom of the staircase releases ADP, disengages from DNA as the ‘seam’ subunit, then repositions to the top to rebind ATP and engage incoming DNA. This process advances translocation by a proposed 1-2 nt per ATP cycle, depending on the number of nucleotides coordinated by each ATPase subunit (30). An alternative inchworm-like mechanism recently proposed for eukaryotic MCM dispenses with strict rotational cycling. Instead, the helicase advances along a 12-nt DNA segment through the planar-to-spiral and spiral-to-planar transitions that pull DNA through the central channel (31–34). Because each of the six MCM subunits coordinates two nucleotides, a complete conformational cycle corresponds to translocation of ∼12 nt and remains consistent with an overall step size of ∼2 nt per ATP hydrolysis event, similar to that proposed for hand-over-hand translocation. Whether either model requires strictly ordered hydrolysis remains debated. Structural studies usually show ATP preferentially binding to adjacent non-seam subunits; however biochemical studies of MCM and the unfoldase ClpX (25, 27, 29, 32) suggest that ATPase-mutant subunits are tolerated, suggesting that hydrolysis can occur with some degree of stochasticity regardless of the dominant translocation mechanism.

Most mechanistic insights into hexameric helicase function derive from structures of single hexamers encircling ssDNA or ssDNA tails of forked substrates, which are generally interpreted as active translocation intermediates (8, 10–13, 17, 19, 21, 35, 36). Capturing helicases bound to duplex DNA has proven considerably more challenging as ring-shaped helicases, such as archaeal and eukaryotic MCMs, tend to slip off duplexes in the presence of ATP or displace one strand during sample preparation (37). Where observed, dsDNA-bound hexameric helicases typically adopt double-hexamer assemblies; in eukaryotes, the origin recognition complex (ORC) is required to load MCM double hexamers onto DNA (18, 38–41). The mechanism of MCM double hexamer formation is debated as single-molecule experiments have suggested sequential loading, in which the first hexamer promotes the loading of the second hexamer. Biochemical data have shown that two distinct engagement events between ORC and DNA promote simultaneous loading of two rings (42–45). How MCM double hexamers transition from an origin-loaded, dsDNA-bound state to an active, unwinding-competent form remains poorly understood, and whether analogous mechanisms operate in minimal self-loading initiator helicase systems remains an open question.

*Staphylococcus aureus* pathogenicity islands (SaPIs) are 14- to 17-kb mobile genetic elements encoding superantigen toxins and other virulence factors (46–48). Although excision has been observed for other pathogenicity islands, SaPIs are unusual in their ability to replicate after excision (46, 48). Each SaPI encodes a primase and a replication initiator protein (Rep) as well as a cognate origin of replication (*ori*) containing six to twelve short repeats (“iterons”) located immediately downstream of the Rep gene (48, 49). Previous studies have suggested that SaPI-encoded Reps function as self-loading homohexameric helicases capable of initiating DNA replication autonomously, combining origin recognition, loading, and melting that in other systems require dedicated accessory factors that in eukaryotes are distributed across dedicated accessory factors including ORC for loading, and Cdc45 and GINS for activation of the MCM complex (34, 50). This independence makes SaPI-encoded Reps attractive systems for dissecting the minimal structural requirements for helicase loading and activation.

Several SaPI-encoded replication initiators (Reps) have been biochemically and/or structurally explored, including those from SaPI1, SaPI2, SaPIbov1, SaPI5 (48, 49, 51, 52), as well as the distantly related Cch and Cch2 helicases from *staphylococcal cassette chromosome* (SCC) elements (53). These Reps share a common biological role but harbor diversified ATPase folds: Reps from SaPI1, SaPI5 and Cch2 belong to superfamily 3 (SF3) (52, 54), while SaPI2, SaPIbov1 Rep and Cch belong to superfamily 6 (SF6) of AAA+ ATPases (53, 54), suggesting that SaPIs may have independently recruited distinct helicase families to drive replication. We previously determined a crystal structure of Cch bound to a non-hydrolyzable ATP analog (53), which revealed a tripartite domain organization shared with archaeal and eukaryotic MCM, comprising an N-terminal domain (NTD) corresponds to the first half of DUF927 that forms an N-terminal ring, a central ATPase domain belonging to SF6 (54) and clade 7 (4), in which the small lid domain is placed such that the sensor II arginine within it acts in *trans* with ATP bound to a neighboring subunit, rather than in *cis*, and a C-terminal winged-helix (WH) domain. In MCM, the WH domain primarily interacts with other proteins (17, 55), whereas in SaPI Reps, it recognizes iteron repeats within the cognate *ori* (48). Notably, whereas MCM engages DNA through pore loop 1 (PL1), pre-sensor 1 β-hairpin (PS1), and helix-2 insertion (H2i) within the ATPase domain, both SaPIbov1 Rep and Cch retain PL1 and PS1, but lack H2i entirely, raising the question of how this subfamily engages DNA in its absence, a question that remained open without a DNA-bound structure of any SaPI-encoded SF6 Rep.

Here we focus on SaPIbov1 Rep, the helicase encoded by and required for the replication of *Staphylococcus aureus* pathogenicity island bovine 1 (SaPIbov1), which shares the tripartite domain organization of Cch. We present the crystal structure of apo SaPIbov1 Rep and cryo-EM structures of a holo complex of SaPIbov1 Rep with AMP-PNP, ADP, and a duplex DNA. Using 3D variability analysis (3DVA) (56), we resolved multiple conformational intermediates revealing how nucleotide-dependent structural asymmetry modulates duplex DNA positioning within the hexameric ring, and show that PL1 and an elongated PS1 constitute the primary DNA-engaging surface in the absence of H2i, with both DNA-interacting motifs sampling distinct configurations with DNA at the base of the staircase. These structures provide the first DNA-bound view of a SF6 SaPI-encoded Rep and offer a structural framework for understanding how this minimal self-loading initiator helicase engages DNA. Additionally, we show that the N-terminus of Rep binds its cognate primase, and we use AlphaFold to suggest how these proteins may recruit additional chromosomally-encoded replication machinery.

## Material and Methods

### SaPIbov1 Rep cloning and protein purification

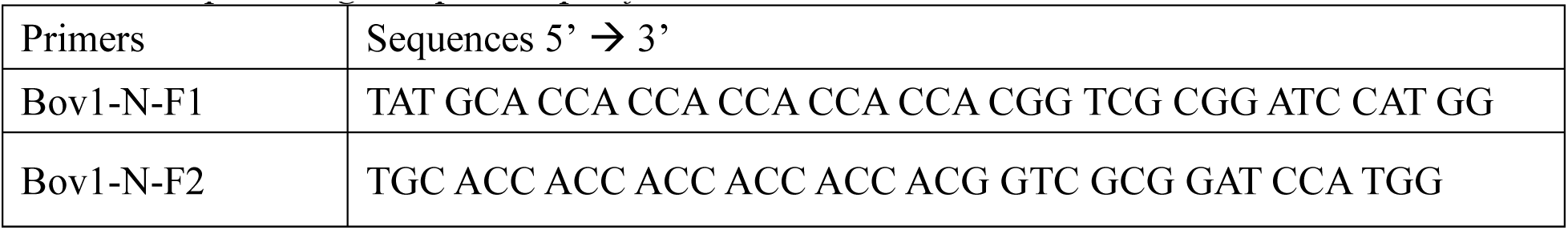
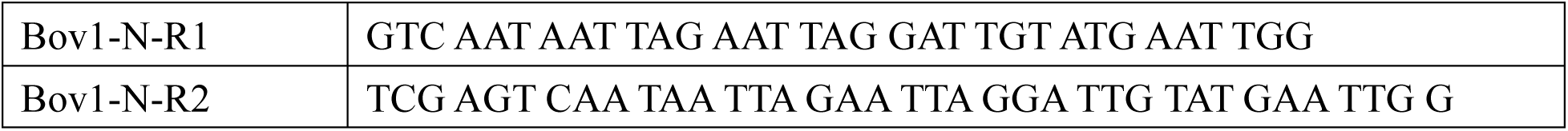

The SaPIbov1 *rep* gene was amplified from strain RN4220 SaPIbov1 *tst*:*tetM* and cloned into plasmid pET-21a into NdeI and XhoI restriction sites using sticky end oligonucleotides listed above. The resulting amino acid sequence carries an extra peptide whose sequence MHHHHHHGRGS is found prior to the initial methionine of the native protein. Native and selenomethionine-labeled protein were expressed and purified as for native and labeled Cch in reference (53), including Ni, heparin and monoS chromatography, except that the final storage buffer was 20 mM tris pH 8.0, 0.5 mM EDTA, 200mM NaCl, 20% glycerol, and 5mM DTT.

### SaPIbov1 Pri-Rep complex cloning and expression

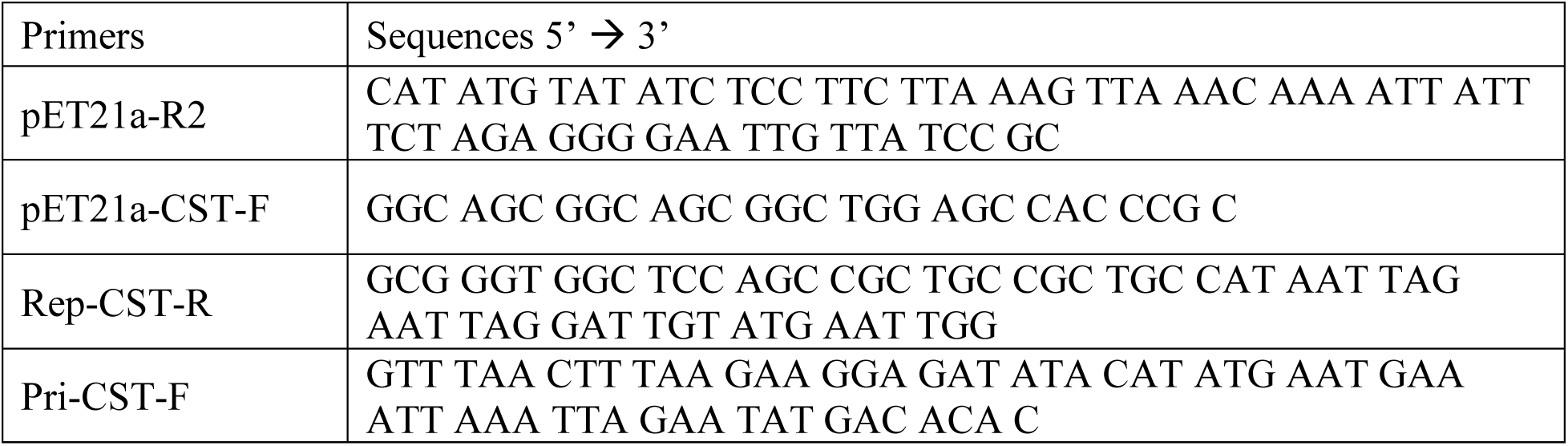

Purified PCR products of a wild-type SaPIbov1 primase (Pri) containing a stop codon, a linker (5’-AAA GGG GCT TTG T-3’), and a wild-type Rep helicase were assembled into a pET21a plasmid using NEBuilder HiFi DNA assembly (New England BioLabs). The assembly of fragments added a flexible linker (5’-GGC AGC GGC AGC GGC-3’) and a Twin-strep-tag (Twin-Strep-tag®) to the C-terminus of Rep.

The Pri-Rep complex was expressed in *E. coli* Rosetta (DE3) pLysS cells. Cells were grown in LB medium with 100 µg/ml ampicillin at 37 ℃ until OD_600_ of 0.6 was reached, then were induced with 0.5mM isopropyl-β-D-1-thiogalactopyranoside (IPTG). After induction, cells were grown overnight at 20℃ and pelleted by centrifugation at 8,000 r.p.m. in an F10S rotor for 10 min. Cell pellets were kept at -80℃ until further use.

### SaPIbov1 Pri-Rep complex purification

Cell pellets were resuspended in lysis buffer (100mM Tris-Cl pH 8, 150 mM NaCl, 1 mM EDTA), lysed by sonication, and clarified by centrifugation at 18,000 r.p.m. in an SS-34 rotor at 4 ℃ for 1h. The filtered supernatant was loaded onto a StrepTrap-HP column (Cytiva) and eluted with elution buffer (100 mM Tris-Cl pH 8, 150 mM NaCl, 1 mM EDTA, 2.5 mM desthiobiotin).

The cleanest peak fractions containing the co-migrating SaPIbov1 Pri-Rep complex were dialyzed against buffer (20mM Tris-Cl pH 8, 100mM NaCl, 0.5 mM EDTA, 5 % glycerol), loaded onto heparin FF 16/10 (Cytiva), and eluted with a linear gradient to buffer containing 20 mM Tris-Cl pH 8, 2 M NaCl, 0.5 mM EDTA, 5 % glycerol. Purified SaPIbov1 Pri and Rep complex was concentrated to ∼16.76 mg/ml and dialyzed overnight at 4℃ against storage buffer (20 mM Tris-Cl pH 8, 200 mM NaCl, 1 mM EDTA, 2 mM DTT, 20 % glycerol). Samples were aliquoted, snap frozen in liquid nitrogen, and stored at -80℃.

### SaPIbov1 Rep crystallization, data collection, structure determination, and refinement

Native and SeMet SaPIbov1 Rep were crystallized using the hanging-drop vapor diffusion method. Although the protein crystallized in the apo-form, single-stranded DNA and a non-hydrolyzable ATP analog were included. The complex was prepared in a total volume of 100 ul, containing: 25 ul of Rep (12 mg/ml in storage buffer; final concentration 3 mg/ml), 25 ul 5x binding buffer (100mM Tris, pH 8, 250mM NaCl, 25% glycerol), poly (dT)_25_ (final protein:DNA ratio 6:1), AMPPNP (final concentration 1mM), MgCl2 (final concentration 1mM) and DTT (final concentration 10mM). Complex was mixed in a 1:1 ratio with mother liquor consisting of 0.1M MES pH6, 25% Jeffamine M-600, 12.5% Ethylene Glycol, 5mM DTT.

X-ray diffraction data for native SaPIbov1 Rep was collected on the GM/CA 23-ID-D beamline at the Advanced Photon Source, Argonne National Laboratory, and the SeMet SaPIbov1 Rep was collected on the GM/CA 23-ID-B beamline. The native crystal diffracted to 2.7 Å whereas a SeMet diffracted to 3.5 Å. All X-ray data were indexed, integrated, and scaled using HKL-2000 software package (57). Both data sets indexed well in R3 space group, but prior to phase determination, the native data set was reindexed with CCP4 to avoid ambiguity in axis assignment issues inherent to this space group. Experimental phases and map were determined using Phenix (58) Autosol with both native and derivative MTZ files. Coot (59) was used for manual model building with iterative refinement using Phenix refine. The final model with R-work and R-free of 0.2651 and 0.2815, respectively, was validated with MolProbity (60) and deposited in the Protein Data Bank. A summary of the data collection and refinement statistics is provided in Extended Data Table 1.

### Sample preparation for cryo-EM

To prepare the DNA-bound SaPIbov1 Pri-Rep complex, 1 mM ATP and 6 mM MgCl_2_ were added to a final volume of 500 µl containing 1.5 mg/ml of purified sample, and the sample was incubated at room temperature for 20 min before being loaded onto a Superose 6 Increase 10/300 GL column (Cytiva) pre-equilibrated with SEC buffer (20 mM Tris, pH 8, 200 mM NaCl). Peak fractions containing the Pri-Rep complex were pooled and concentrated to 0.3 mg/ml using an Amicon Ultra centrifugal filter concentrator (50 kDa, MilliporeSigma). A 15 µl aliquot of the sample was then incubated at room temperature for 10 min in the presence of 0.5 µM 25 nt ssDNA (5’-CAA GCT TGT TTT TCC ATA GGC TCC G -3’), 3 mM AMP-PNP, 3 mM ADP, and 6 mM MgCl_2_, in a final volume of 20 µl.

### Grid preparation for cryo-EM

Freshly glow-discharged C-flat 1.2/1.3 400 mesh holey carbon grid was loaded into a Vitrobot^TM^ mark IV (FEI) chamber operated at 18 ℃ and 100 % relative humidity. 3 µl of sample was applied to the grid, blotted for 5.5 s with blot force set to 0, and plunge-frozen into liquid ethane.

### Data collection for cryo-EM

Data were collected on a Titan Krios G3i electron microscope (Thermo Fisher Scientific) operating at 300kV and equipped with a Gatan K3 direct detection camera. Data collection was performed at the Advanced Electron Microscopy Facility, the University of Chicago. Each movie consisted of 50 frames with a total exposure of approximately ∼50 electrons/Å2. The defocus range was set at – 1.0 to – 2.5 µm. Additional acquisition parameters are provided in Extended Data Table 2.

### Image processing

All image processing was performed in cryoSPARC (v.4.6.0) (61) and summarized in (Supplementary Figure 1). Movie stacks were motion corrected using patch motion correction, and contrast transfer function (CTF) parameters were estimated using patch based CTF estimation. Micrographs with significant non-vitreous ice contamination or poor motion correction and CTF parameters were excluded.

Initial particle picking was performed using the blob particle picker, followed by two-dimensional (2D) classifications. After a second round of 2D classification, selected classes showing intact complexes with secondary structures were used as templates for automated template-based particle picking. False-positive, aggregated, or poorly defined particles were filtered out after iterative rounds of 2D classification. The remaining 607,797 particles were used to train a Topaz (v.0.2.4)(62) model, which was then applied to the micrographs for refined particle picking.

Two three-dimensional (3D) ab initio models were generated with C1 symmetry. The class representing the intact DNA-bound complex was refined using non-uniform refinement. Ten 3D classes were generated, followed by five classes exhibiting well-defined features were selected for heterogeneous refinement. A total of 225,203 particles from two representative classes displaying two distinct modes of intersubunit arrangements of the two highly dynamic adjacent subunits were selected for reference-based motion correction. After particle cleaning, 217,928 good particles were used for final homogeneous refinement, resulting in a consensus 3D reconstruction at an overall resolution of 3.06Å, as determined by the Fourier shell correlation (FSC) based on the 0.143 criterion. Local resolution was estimated using cryoSPARC (61).

Conformational heterogeneity was further investigated using 3D variability analysis (3DVA) (56) with four variability modes and a filter resolution of 5Å. Latent coordinate distribution displayed a spherical pattern, consistent with continuous conformational variability. For each variability mode, 10 frames were sampled along the principal component using 3DVA ‘simple mode’, and the first and last frames (frame 0 and 19) were selected as representative conformational endpoints. These maps, together with particle subsets generated from the 3DVA ‘intermediate mode’ were subjected to non-uniform refinement. Movies depicting conformational transitions were generated within UCSF chimera (v.1.17.3) (63).

### Model building, refinement and validation

The crystal structure of apo SaPIbov1 Rep helicase was rigid-body fit into the cryo-EM map using ChimeraX (v.1.4) (64). A partially duplexed B-form DNA with 3’ ssDNA tail was generated using AlphaFold3 (65) and positioned according to the cryo-EM map using ChimeraX (64). Overlapping regions were manually corrected and adjusted using ISOLDE (v.1.4) (66). Manual model building and refinement were performed iteratively using COOT (v.0.9.8.8) (59), ISOLDE (66), and Phenix (v.1.21) (67) real-space refinement. Geometry restraints for nucleotide binding sites were applied based on AMP-PNP and ADP and Mg^2+^ binding models. Model quality was assessed using MolProbity. Figures and structural analyses were generated using ChimeraX (64) and PyMOL (v.3.1.3) (68).

## Results

### Apo SaPIbov1 Rep forms a symmetric homohexameric ring

Our crystal structure of SaPIbov1 Rep provides a view of the apo form. The structure was determined at 2.7Å resolution and contains two molecules per asymmetric unit. A non-crystallographic two-fold symmetry axis superimposed on a crystallographic three-fold axis generates a planar hexameric ring (Figure 1). The narrowest regions within the central channel of the NTD, ATPase, and C-terminal WH domain measure ∼23Å, 35Å and 16Å, respectively, sufficient to accommodate both ssDNA and dsDNA, with additional space permitting conformational breathing (Figure 1A). AlphaFold3 models predict ssDNA and dsDNA threading through the SaPIbov1 Rep helicase (Supplementary Figure 2).

**Figure 1.**
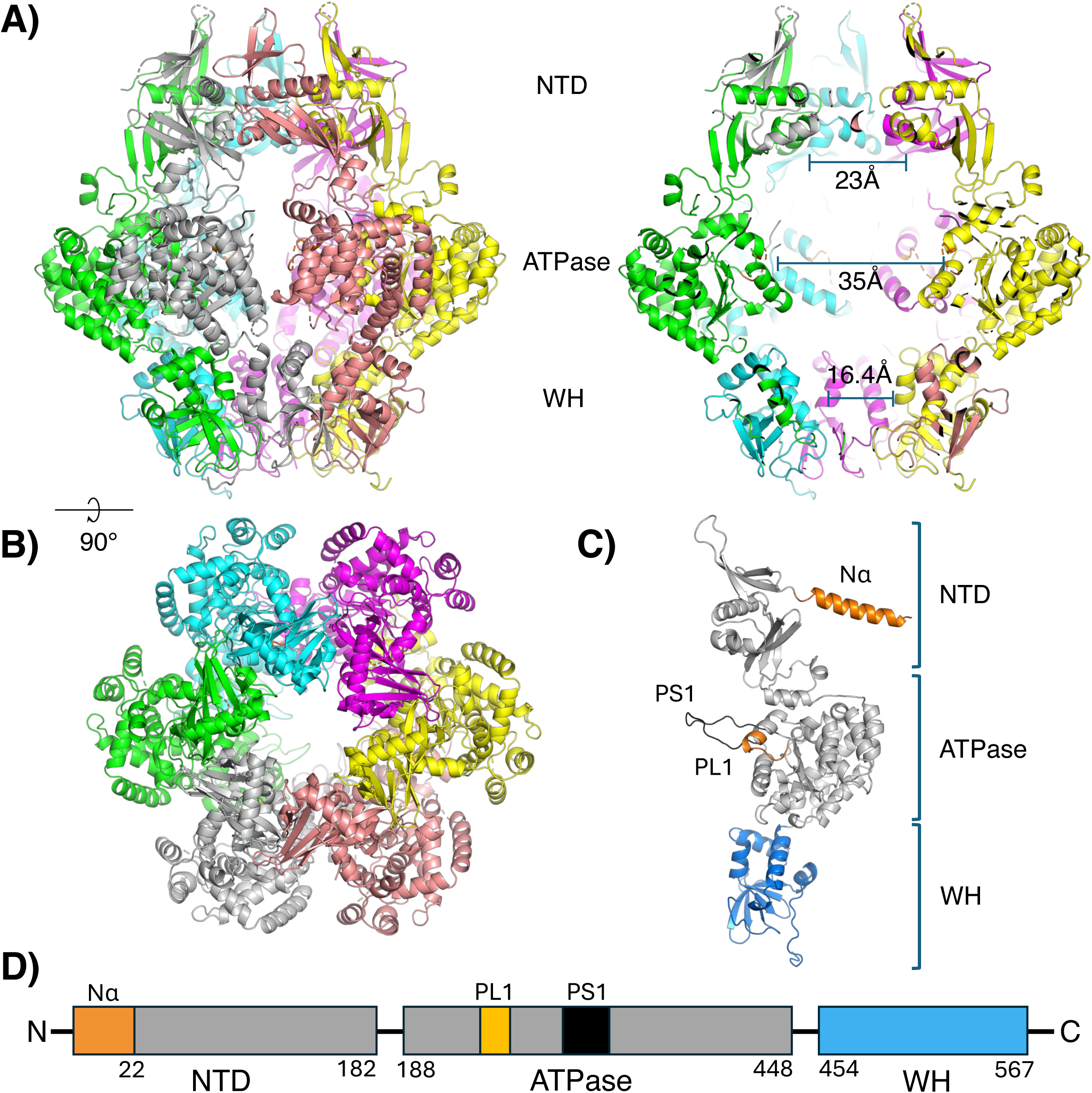
Crystal structure of apo SaPIbov1 Rep. A. Biological assembly of SaPIbov1 Rep in the apo state, shown in side (left) and sliced (right) views. The asymmetric unit contains two adjacent subunits. B. Ribbon diagram shown in top view. C. Ribbon diagram of a SaPIbov1 Rep monomer predicted by AlphaFold2, colored as in (D). D. Linear schematic of SaPIbov1 Rep domain organization. Nα: the first helical segment (orange, residues between 1-22); NTD: N-terminal domain from residues 1-182; ATPase: residues between 188-448); PL1: pore-loop 1 (pale orange, residues between 257-265); PS1: presensor 1 β hairpin (black, residues between 301-318); WH: C-terminal WH domain (blue, residues 454-567).

The overall architecture of SaPIbov1 Rep closely resembles that of the Cch helicase (PDB 5DGK), despite a low sequence percent identity of ∼16.2%. Both structures adopt a planar hexameric ring geometry in the absence of DNA (Supplementary Figure 3). The NTD consists of two anti-parallel β-sheets on the exterior of the ring, with α-helices lining the interior (Supplementary Figure 4A). Interfaces between NTDs are stabilized primarily by hydrophobic interactions among the helices. Although both SaPIbov1 and Cch lack the zinc-binding motif conserved in MCM N-terminal domains, they feature positively charged Lys and hydrophobic Ile residues near the tip of a β-hairpin that protrudes from the top of the hexameric ring. These residues are structurally reminiscent of Arg195 and Leu209 in the double hexameric MCM complex (41) (Supplementary Figure 4B), which have been implicated in stabilizing the separated DNA bases formed during the initial melting of the DNA duplex.

Two DNA-binding motifs from the central ATPase domain of each subunit line the central channel. The first is the pre-sensor 1 β-hairpin (PS1) insertion, between α3 and β4, a DNA-binding motif that is shared across replicative hexameric helicases in the PS1 insert superclade (clades 4-7) (2, 7). The second pore-lining motif is located between β2 and *a*2; this region is typically referred to as pore loop 1 (PL1) in classic protein-remodeling AAA+ proteases in clade 3, such as Vps4, katanin, ClpB, and Hsp104 (7). Unlike the canonical PL1 in clade 3, the PL1 region found in SaPIbov1, Cch, and MCM helicases lacks both the α-insert and the conserved aromatic residue involved in substrate recognition (Supplementary Figure 5). Previous studies have noted that the PL1 region in MCMs adopts either a horseshoe or helical conformation at loose or tight subunit interfaces, respectively (14, 20). Although the position of the lid subdomain relative to the ATPase places both SaPIbov1 and Cch ATPases in clade 7, along with MCM, they lack the helix-2 insertion (H2i) motif that is otherwise conserved within clade 7 (Figure 1 and Supplementary Figure 5). Instead, both SaPIbov1 Rep and Cch possess an elongated PS1 relative to MCM, suggesting that this extension may have evolved to functionally compensate for the absence of H2i (or that H2i evolved separately to enhance DNA binding). Both DNA-interacting motifs of SaPIbov1 Rep exhibit elevated B-factors in the absence of DNA (Supplementary Figure 3).

The C-terminal WH domain is primarily composed of a winged-helix-turn-helix motif (Figure 1D and Supplementary Figure 6). Unlike the NTD, the WH domain’s structure and function appear conserved among other SaPI-encoded Reps. Previous biochemical and structural studies have shown that this domain binds short DNA repeats (iterons) at the origin (48, 49). Our previous structural study of Cch indicated that the dsDNA-binding region is occluded by the adjacent subunit in the hexameric ring (53), requiring substantial conformational rearrangement for exposure and dsDNA binding. AlphaFold 3 predictions of the isolated WH domain of SaPIbov1 Rep also show the dsDNA binding region buried by the adjacent subunit in the ring (Supplementary Figure 6D). AlphaFold3 predicts that the WH domain uses an unusual DNA-binding mode that places the beta strands rather than a helix into the major groove (Supplementary Figures 6 and 7). A similar binding mode was reported for TcpK, which performs a similar role – it binds iterons found in the replication origin of a mobile genetic element (a conjugative plasmid) (Supplementary Figure 7F). Despite only ∼21% sequence identity, TcpK was the top experimental structure match found by Foldseek (69). Furthermore, both SaPIbov1 and TcpK recognize similar iterons sequences.

While structural comparison of individual domains in SaPIbov1 Rep and Cch highlights overall interdomain stabilization in a planar hexameric ring architecture without DNA, differences in nucleotide binding state at the ATPase subunit interfaces result in distinct ATPase domain arrangements within the tier. In Cch, AMP-PNP, Mg^2+^, and three water molecules are stabilized in a canonical ATP-binding configuration, with *trans*-acting arginine finger and sensor II residues coordinating the ɣ phosphate, creating tight subunit interfaces. In contrast, apo SaPIbov1 Rep lacks nucleotide occupancy at the active sites, resulting in looser intersubunit contacts and likely contributing to higher average B-factors in the ATPase domains (Supplementary Figure 3A). Moreover, ATP-analog stabilization in Cch positions ATPase domains around the ring such that the PS1 hairpin from one protomer partially aligns above the PL1 of the neighboring subunit, further enhancing intersubunit stabilization.

### Substrate binding induces an asymmetric staircase architecture

Our initial cryo-EM efforts with DNA-bound SaPIbov1 Rep alone were hindered by severe orientation bias. To overcome this limitation, we co-purified Rep bearing a C-terminal twin strep tag with its upstream SaPIbov1 Primase (Pri), which formed a semi-stable complex in solution (Supplementary Figure 8D). AlphaFold3 modeling predicts high confidence interactions between the N-terminal helix of Rep and the C-terminal domain of Pri (Supplementary Figure 8). Consistently, truncation of the N-terminal helix abolished complex formation in solution (Supplementary Figure 8D). Notably, this structural element is absent in the SCC-encoded Cch. This divergence correlates with the fact that Cch-encoding SCC elements do not carry an identifiable primase gene, and suggests that these SCCs and SaPIs have evolved different strategies for priming DNA synthesis.

We reconstituted the Pri-Rep complex with a 25-nt ssDNA substrate as well as AMP-PNP, ADP and Mg^2+^ to allow different nucleotide states without actual hydrolysis. The inclusion of Pri substantially mitigated orientation preference, yielding a consensus reconstruction of the Rep hexamer at a global resolution of 3.06Å; however, density corresponding to Pri and the N-terminal helix of Rep was absent during data processing and in the final map (Supplementary Figure 1), likely reflecting dissociation of the primase during grid preparation, a random orientation relative to the helicase, and/or naturally sub-stoichiometric occupancy (note only 1 copy of primase is needed per helicase to prime lagging strand synthesis).

In contrast to the SaPI1 Rep cryo-EM structure, in which the addition of duplex DNA resulted in binding of a single-stranded penta-nucleotide substrate within the central channel (52), the SaPIbov1 Rep map revealed partially duplexed DNA in the central channel even though the substrate was intended to capture ssDNA translocation intermediates. Eight nt at the 5’ end of the substrate were fortuitously self-complementary, and despite a calculated melting temperature of ∼12.2 ℃, they are clearly base-paired and bound along the inner lining of the ATPase domains, with a periodicity of 2 bp per interacting subunit (Figure 2 and Supplementary Figure 9B). Partially resolved density for one 3’ ssDNA tail extends toward the NTD entry and threads through the central pore, where it is stabilized by Arg104 residues encircling the narrow pore (Supplementary Figure 4A and 9A). In contrast, density for the opposing 3’ ssDNA extension toward the C-terminal WH domain exit is limited to the first two nucleotides (Supplementary Figure 9A and B), reflecting weaker interactions near the C-terminal WH domain.

**Figure 2.**
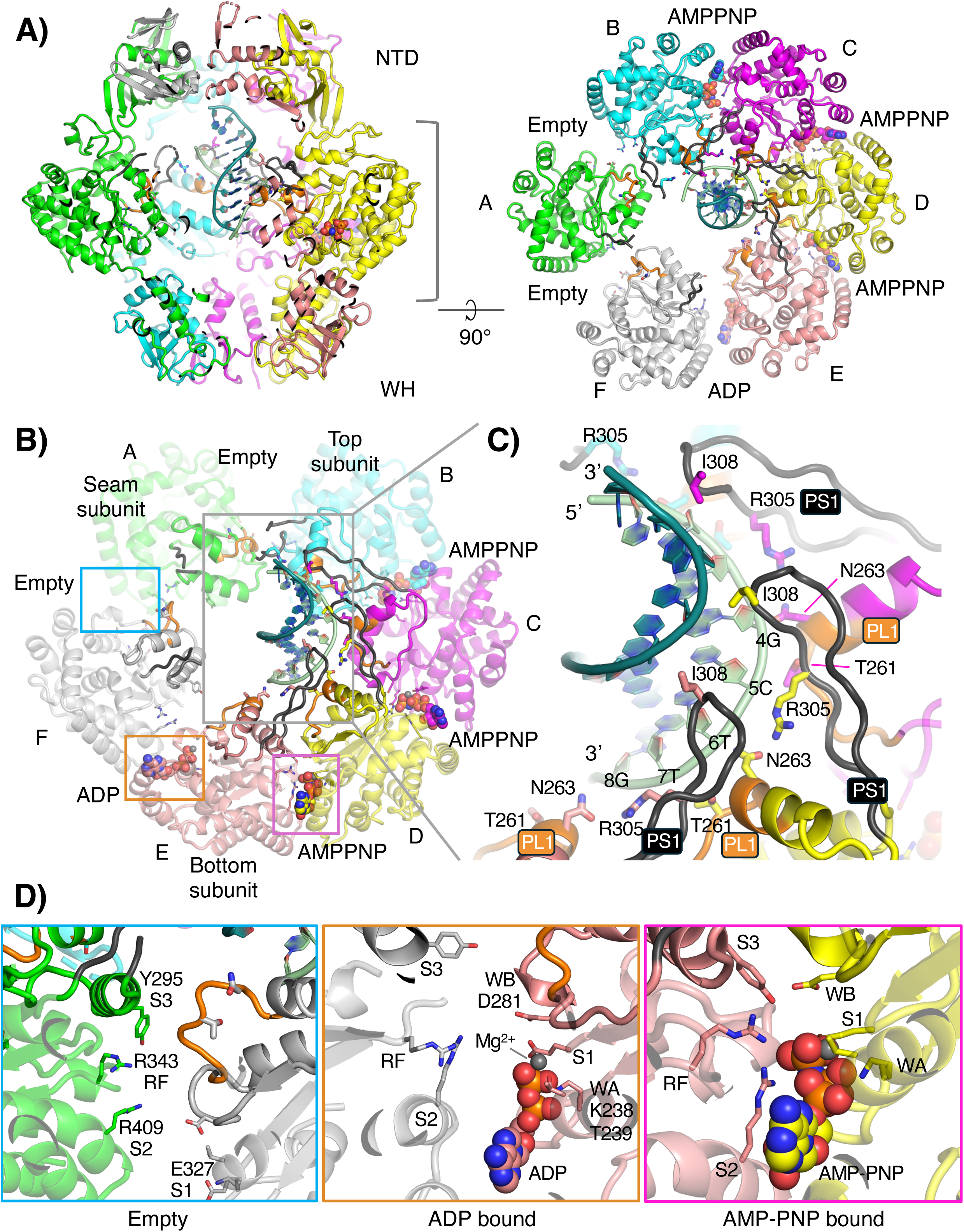
Cryo-EM structure of SaPIbov1 Rep in the dsDNA- and nucleotide-bound state. A. Ribbon diagrams of SaPIbov1 Rep shown in sliced side (left) and sliced top (right) views, highlighting dsDNA, AMPPNP, and ADP bound within the central channel of the ATPase tier. The color scheme corresponds to the Figure 1 – each subunit is a different color, except for PL1 (orange) and PS1 (black). The gray bracket indicates the range of focal plane for the top view. B. ATPase domains are oriented such that subunit B at the top of the staircase is positioned at the top, with the descending subunits arranged clockwise and the bottom subunit (E) positioned at the bottom. C. A close-up view of dsDNA binding by the pore-loop 1 (PL1; orange) and pre-sensor 1 β-hairpin (PS1; black) motifs within the central channel, oriented as in panel B (gray box). The leading strand is colored light green and the lagging strand is colored dark teal. Key PS1 residues (R305 and I308) and PL1 residues (T261 and N263) are shown as sticks. D. Close-up views of the intersubunit nucleotide binding sites, indicated as in panel B (empty site, blue box; ADP-bound site, orange box; AMP-PNP bound site, magenta box). Conserved residues involved in nucleotide binding and intersubunit interactions are labeled and shown as sticks.

The ATPase domains of the holo structure adopt a lock washer or spiral staircase geometry. AMP-PNP is present at the B-C, C-D, and D-E subunit interfaces, ADP occupies the E-F interface (Figure 2A and Supplementary Figure 9C), and the F-A and A-B interfaces are empty. Subunit A, which is flanked by “empty” interfaces, is the least well ordered and lies at the “seam”, or the crack in the lock washer. At AMP-PNP bound interfaces, the ATPase domains adopt configurations similar to those observed in Cch, in which the sensor III residue Tyr295 forms a H-bond with the sensor I residue Glu327 of the adjacent subunit (Figure 2D and Supplementary Figure 3B). This interaction further strengthens the tight intersubunit interface and positions the PS1 hairpin of one subunit above the PL1 motif of its neighboring subunit (Figure 2B). Nucleotide binding correlates with DNA binding. The asymmetry in DNA binding arises primarily from extensive engagement of the partial duplex region by the PS1 hairpin, PL1 motifs, or both of subunits B, C, D, and E (Figure 2). Coordinated engagement of the leading strand by these motifs drives the formation of a right-handed spiral staircase arrangement of the ATPase domains spanning subunits B, C, D, and E. The staircase descends in the order B→C→D→E, with AMP-PNP occupying the upper interfaces (B-C, C-D and D-E) and ADP at the base subunit interface E-F (Figure 2B). Specifically, the PS1 hairpins from subunits C, D, and E align along the leading strand, where the guanidinium groups of Arg305 are positioned in close H-bonding distance with the phosphates of T-9, T-7 and C-5, respectively (Figure 2C). Ile308 at the tip of the PS1 hairpin lies within the major groove, making non-specific van der Waals contacts with bases at positions n-2 and n-3 (relative to the Arg305 contact from the same subunit) (Figure 2C). Concurrently, Thr261 and Asn263 from the PL1 motif of the adjacent subunit coordinate the phosphate at the n-1 position (Figure 2C).

The resulting spiral staircase configuration generates asymmetric DNA engagement among subunits. Subunits C and D, positioned in the middle of the staircase and with full AMP-PNP occupancy at both interfaces, exhibit full engagement of the DNA via their PS1 and PL1 motifs (Figure 2B). In contrast, subunit B at the top of the staircase, with only one AMP-PNP bound interface (B-C), displays a partially disordered PS1 hairpin. Here, the duplex DNA is stabilized below the PS1 tip, offering no structural platform to support Ile308 engagement (Figure 2B). Similarly, subunit E at the base of the staircase, bearing one AMP-PNP and one ADP interface, shows partial disengagement of its PL1 motif from the substrate; the duplex DNA region lies above its contact point and the ssDNA extension falls outside of the contact distance, and only Asn263 remains within H-bonding distance (Figure 2C). The duplex DNA in the lower part of the staircase adopts an intermediate conformation with an A-B DNA index (ABI) (70) of 0.49, whereas toward the top of the staircase the duplex transitions to a more unwound state with an ABI of 0.15, indicative of increase in A-form character. This transition along the staircase likely reflects the strong DNA grip exerted by subunit C and D through their fully engaged PS1 and PL1 motifs including Ile308-mediated contacts.

Collectively, these structural features and the low Tm of our 8bp duplex indicate that SaPIbov1 Rep may preferentially bind duplex DNA over ssDNA within its central channel, or that it preferentially binds the ssDNA-dsDNA junction. Notably, we observed no evidence of strand separation within the central channel, such as localized melting or strand extrusion. The short duplex substrate used here under non-hydrolyzing conditions likely prevented progression beyond early translocation, perhaps trapping the equivalent of half of the double hexamer that MCM initially forms on duplex DNA before transitioning into a full unwinding mode.

### Seam subunits exhibit continuous conformational sampling in a closed ring

Previous cryo-EM studies of other SaPI-encoded Reps that belong to the SF3 group, such as SaPI1 and SaPI5, reported high conformational variability of the CTD relative to the ATPase domain, and its important role in ATPase activity (52). In contrast, SaPIbov1 Rep exhibits limited CTD motion in our reconstructions (Supplementary Figure 1B). Instead, the most prominent variability in the consensus map is localized to the ATPase domain of the ‘seam’ subunits A and F, despite being under the non-hydrolyzing conditions (Supplementary Figure 1B). These subunits are positioned distal to the DNA and exhibit substantially higher conformational heterogeneity compared to the nucleotide- and DNA-engaged subunits, consistent with the absence of stable nucleotide occupancy at their interfaces. Although ATPase domains of subunits A and F display high B-factors, the helicase maintains a closed hexameric ring, most likely due to the rings formed by the N- and C-terminal domains. The weak or poorly defined density for these ATPase domains in the consensus reconstruction likely reflects averaging of transient conformational states sampled by the seam subunits during refinement, including possible interactions with neighboring subunits and DNA. Together, the gradient in nucleotide occupancy across the staircase, the asymmetric DNA engagement within the ring, and the dynamic nature of the seam subunits suggest that the helicase was captured in a state consistent with continuous translocation-like motion, even under non-hydrolyzing conditions.

To further decouple the conformational heterogeneity masked in the consensus map, we performed three-dimensional variability analysis (3DVA) (56) and analyzed the top three principal components (PCs) (Supplementary Figure 1C and D). For each PC, ten frames were reconstructed from equally spaced positions along the linear trajectory defined by the latent reaction coordinate distributions. The spherical distribution of coordinates within the latent dimensions further supports the presence of continuous conformational heterogeneity in this system. Representative conformations corresponding to the extrema of each variability mode were modeled using the first and last frames of each trajectory.

All PC maps revealed substantial ranges of motion of the seam subunits within the ring. Along the trajectory of the first two variability components, snapshots from frame 0 to 9 show that subunits A and F appear to undergo progressive opening of the A-F interface (Figure 3 and Supplementary movies 1 and 2). In contrast, the third component reveals concerted motion of subunits A and F toward the subunit B, accompanied by a slight upward displacement of subunit A along the spiral staircase toward the subunit B (Figure 3 and Supplementary movie 3). These global motions of the seam subunits likely facilitate transient interactions with adjacent subunits to maintain an overall integrity of the closed ring.

**Figure 3.**
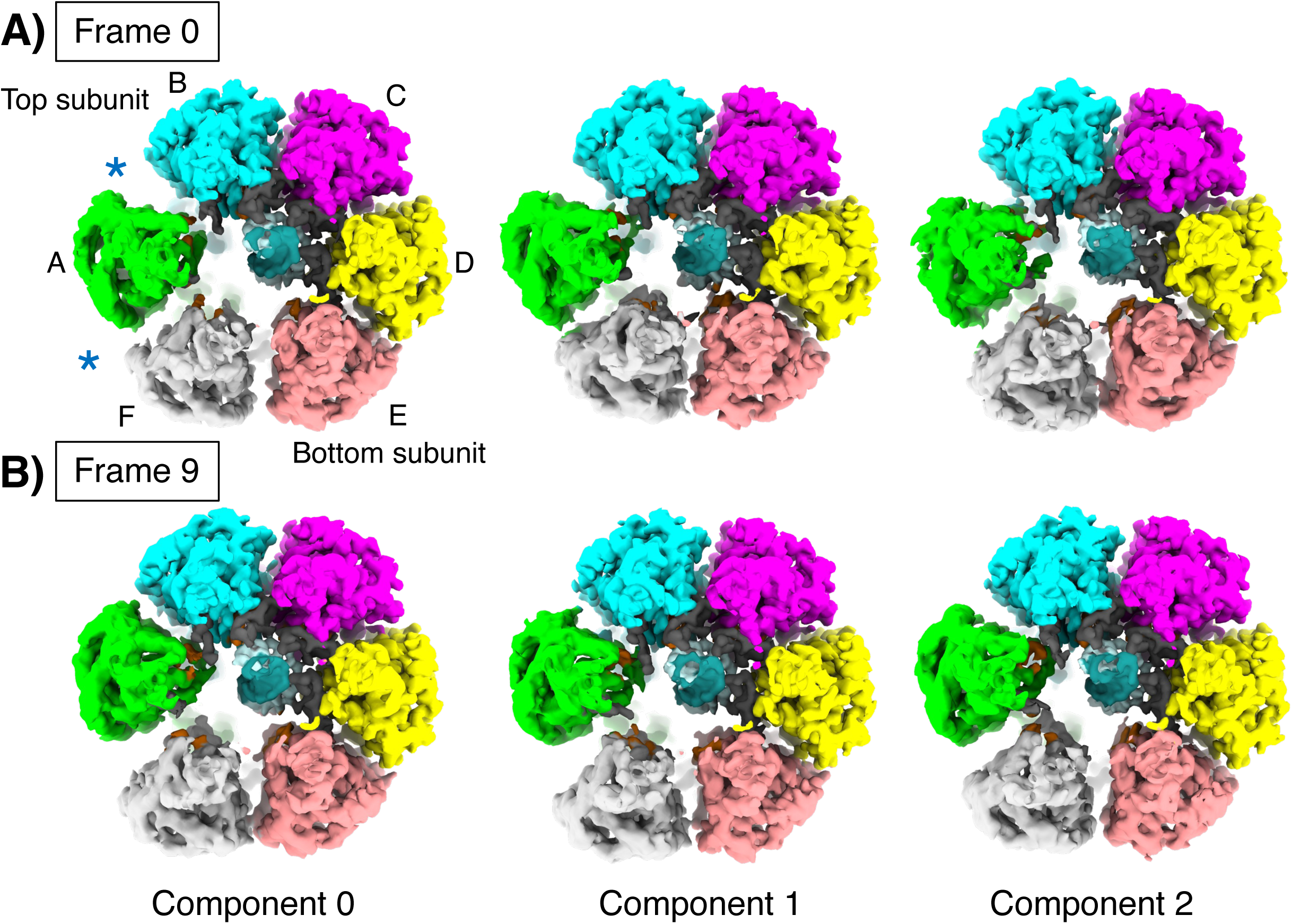
Conformational sampling of seam subunits with neighboring subunits and DNA within the closed ring. A. Three principal components (component 0, left; 1,middle; and 2, right) were derived using 3-D variability analysis (3DVA)(56). For each component’s trajectory, two extrema along the reaction coordinates were used to generate first and last frames (frame 0 and 9). Reconstructed maps of the first frame for each component are shown from the same viewing angle as in Fig. 2A. Subunits are labeled A-F using the same coloring scheme as in Fig 2. Duplex DNA is colored pale green and teal within the central channel. Blue asterisks mark seam interfaces. B. Similar views of the three principal components shown in panel A. Reconstructed maps of the last frame for each component.

### PS1 hairpins in seam subunits transiently engage DNA during conformational sampling

The highly ordered PS1 hairpins in subunits C, D, and E primarily bind the leading strand in coordination with the PL1 motif and contribute to asymmetric DNA engagement within the ring’s spiral staircase (Figure 2). In the consensus map, the PS1 hairpin of subunit B, located at the top of the staircase, is partially disordered (residues 307-312), and the PS1 hairpins in seam subunits A and F are also disordered (residues 303-314), preventing reliable atomic modeling of these regions. Given our previous observations that seam subunits engage in transient conformational interactions with adjacent subunits, we examined the PC maps for potential interactions of the PS1 hairpins in these regions, focusing on seam subunits A and F. At lower contour level, the PC maps revealed that these PS1 hairpins exhibit varying degrees of partial density stabilization as the seam subunits sample the global range of conformational motions within the ring. The PS1 hairpin of subunit A, with unoccupied interfaces on both sides, exhibits greater conformational disorder than that of subunit F, where the E-F interface is partially stabilized by ADP (Figures 2 and 4).

**Figure 4.**
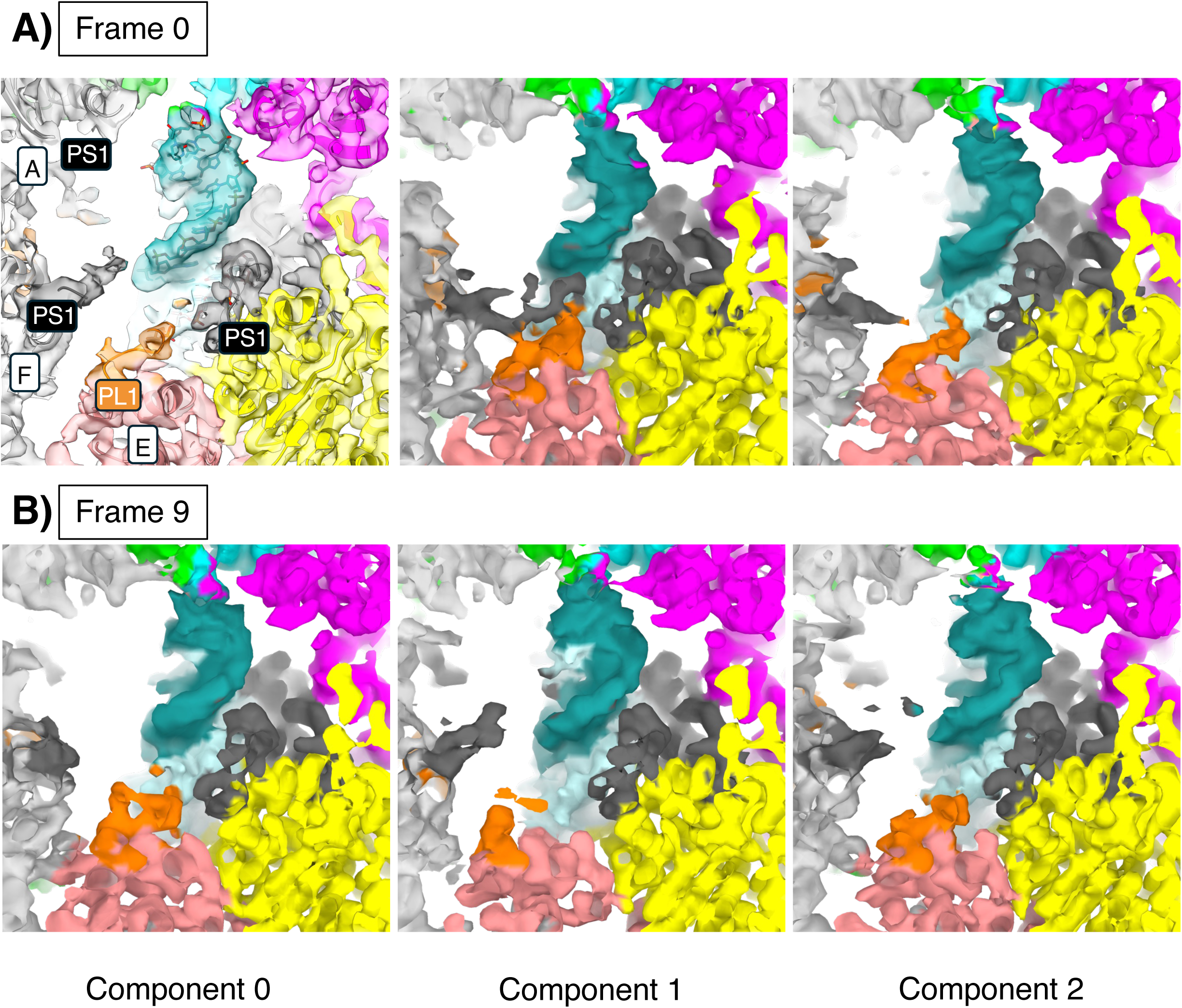
PS1 of seam subunits. A. Close-up comparison of component maps (component 0, left; 1, middle; 2, right) of the first frame using the same coloring scheme as in Fig. 2. All maps were contoured at 0.03 to illustrate relative positional difference in PS1 (black) in stabilizing duple DNA. B. Similar views of the three principal components shown in panel A. Reconstructed maps of the last frame for each component.

The PS1 hairpin of subunit F adopts at least two discrete states: partially ordered through DNA engagement or fully disordered (Figure 4). Among the PC maps showing partial ordering, the PS1 hairpin of subunit F adopts two different positional engagements with DNA: one projecting upward toward the middle of the staircase to transiently bind the complementary strand backbone, and another moving downward into the DNA groove. The latter resembles the PS1 conformation adopted by subunits C,D, and E in the upper staircase (Figure 5). These observations indicate that seam subunits not only interact with their neighboring subunits but also transiently engage DNA during translocation, raising the question of what structural features gate these two PS1-F positional states.

**Figure 5.**
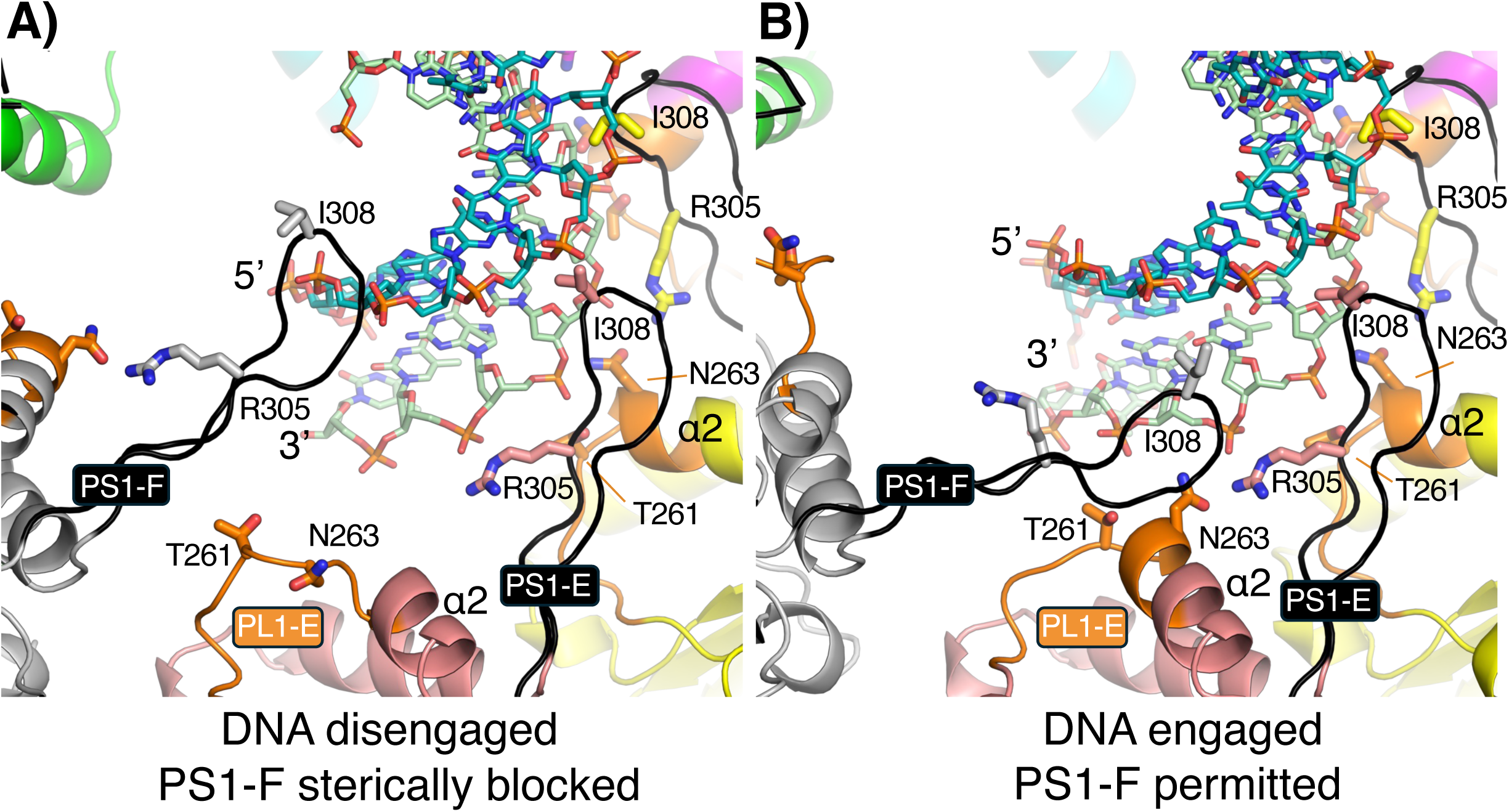
Conformational gating of PL1-E in DNA capture at the base of the staircase. A. A close-up view of the PL1 motif of subunit E (PL1-E) and the PS1 motif of subunit F (PS1-F) at the base of the staircase, showing PL1-E disengaged conformation. Partially destabilized PL1-E helix sterically blocks PS1-F from adopting the DNA engaged state. B. Similar view as in panel A, showing PL1-E engaged with DNA, which permits PS1-F to engage within the DNA groove, extending the descending staircase conformation.

### PL1 conformational switch coordinates PS1 hairpin of neighboring seam subunit for discrete DNA capturing

At the base of the staircase, we also observed conformational switching of the PL1 motif in subunit E between horseshoe and helical conformations. These PL1-E conformational changes appear to correlate with transitions between DNA-engaged and DNA-disengaged PL1-E states, as well as with the changes in the PS1-F positional states described above. In the DNA-disengaged state, PL1-E adopts a horseshoe conformation in which Asn263 undergoes a complete peptide rotamer rearrangement, partially destabilizing the α2 helix and converting it into a flexible loop that swings out of H-bonding distance to the DNA (Figure 5). Conversely, in the helical PL1-E conformation, Asn263 is positioned within H-bonding distance of Arg305 of PS1-E, and additional density for the DNA backbone appears at low contour level within the H-bonding distance of Thr261 and Asn263. These weak and diffused additional densities at the bottom of the staircase may reflect multiple positional states of the duplex DNA, suggesting that DNA engagement by PL1-E is transient in the presence of ADP at the subunit interface.

PS1-B at the top of the staircase exhibits an inverse relationship with PL1-E conformational states at the bottom of the staircase (Figure 3). When PL1-E adopts the DNA-disengaged horseshoe conformation, PS1-B is partially ordered and in close proximity to an additional duplex density observed just above the top of the staircase. In contrast, when PL1-E adopts the DNA-engaged helical conformation, this additional density is no longer observed, and PS1-B becomes disordered. These observations reflect a coupling between PL1-E conformational states, PS1-B ordering, and local DNA positioning at the top of the staircase.

Together, these observations suggest that SaPIbov1 Rep adopts two major coordinated conformational states involving concerted rearrangements of the subunits at the top and bottom of the staircase in response to alternative duplex positioning states within the channel. Conformational switching of PL1 may further gate the next translocation step by controlling the positioning of the PS1 hairpin in the adjacent seam subunit F. The downward insertion of PS1-F into the DNA groove, resembling the position of the bottom subunit of the staircase, is only structurally permissible when PL1 of subunit E is engaged; a disengaged PL1 conformation would introduce steric clash and preclude proper PS1 insertion into the DNA groove. Conversely, when PL1-E disengages, PS1-F is sterically displaced out of the groove. This reciprocal relationship between PL1-E engagement state and PS1-F positioning suggest that PL1 at the base of the staircase acts as a primary structural gate that selectively permits PS1-F insertion as the new bottom subunit only when the DNA is properly captured at the base.

Overall, the dynamic nature of the SaPIbov1 Rep helicase, which undergoes various conformational changes in response to duplex binding including ring opening and closing, PL1 and PS1 mediated DNA binding, and asymmetric subunit repositioning, likely represent intrinsic intermediates of the helicase captured along the DNA translocation pathway.

### Primase is predicted to recruit a host polymerase

The fact that Rep and Primase are the only proteins encoded by SaPIbov1 that are required for its replication implies that they must recruit host-encoded machinery. We found that AlphaFold confidently predicts an interaction between Primase and the N-terminal exonuclease domain of DNA Pol III alpha subunit (71) (Supplementary Figure 8E). It is currently unknown what other replication machinery is commandeered by the replicating SaPI and how it is organized – a question that is particularly interesting because the SaPI helicase translocates along the leading strand, whereas the helicase that drives the chromosomal replication fork translocates along the lagging strand.

## Discussion

Structural comparison of SaPIbov1 Rep with its closest structural homolog Cch and with MCM helicases reveals how conserved AAA+ architecture may have diversified to support distinct replication strategies. Despite low sequence identity, both SaPIbov1 Rep and Cch conspicuously lack H2i, a motif otherwise conserved within clade 7 that, in MCM helicases, contributes a conserved aromatic residue from H2i to a composite “aromatic wedge” formed around the DNA and promotes duplex destabilization (8, 14, 20, 40). Structures of the papillomavirus E1 helicase and SV40 large T-antigen, both SF3 helicases structurally distinct from MCM, show PS1 binding one strand of DNA via a conserved lysine and a subsequent aromatic histidine residue that stacks on DNA sugars, with striking similarity to the MCM H2i aromatic wedge (Figure 6) (20). In both SaPIbov1 Rep and Cch, these conserved residues are missing altogether. Instead, an elongated PS1 motif in these proteins appears to compensate for the absence of both the H2i aromatic wedge and the PS1 hairpin-tip contacts. Rather than contacting DNA solely through its hairpin tip, Arg305 of SaPIbov1 Rep, near the base of PS1, coordinates the DNA backbone, while Ile308 at the PS1 tip makes non-specific van der Waals contacts with bases at positions n-2 and n-3 relative to the Arg305 contact (Figure 6A). This arrangement is shifted relative to MCM, where the aromatic wedge of H2i engages bases at positions n-0 and n-1 relative to a lysine equivalent backbone contact. Thus, whereas the MCM aromatic wedge residue is positioned adjacent to the primary backbone-coordinating residue, the DNA-contacting residues in SaPIbov1 Rep are positioned further apart, allowing the elongated PS1 to engage the DNA backbone along an extended surface.

**Figure 6.**
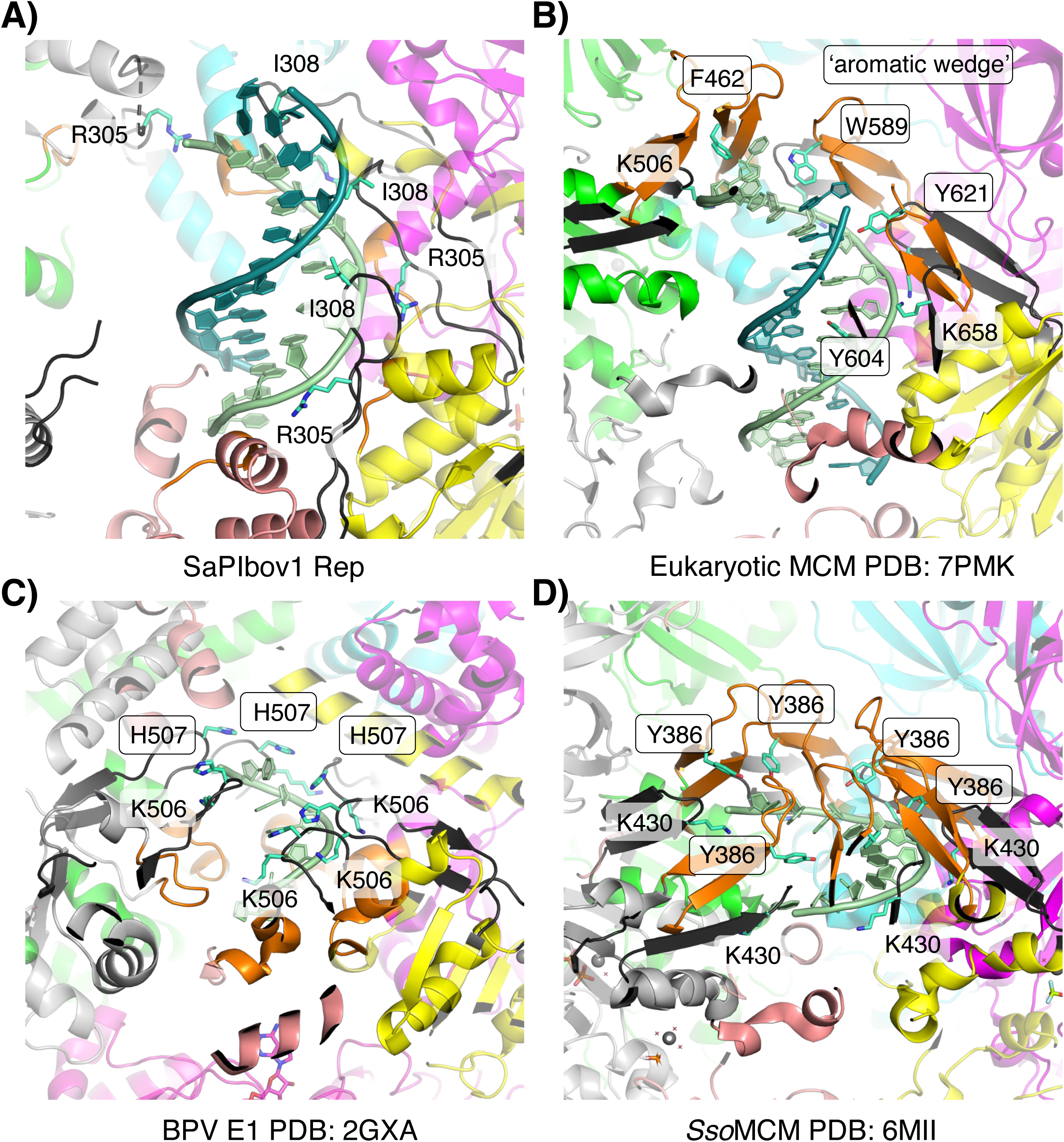
Comparison of DNA binding residues and conserved aromatic wedge residues in ATPases from archaea, eukaryotes, and bovine papillomavirus. A. Cryo-EM structure of SaPIbov1 Rep encircling dsDNA. Key DNA binding residues (R305 and I308) are shown as sticks (cyan). The same coloring scheme is used in panels B-D. Structures are aligned based on SaPIbov1 Rep ATPase domains. B. Cryo-EM structure of eukaryotic MCM (PDB 7PMK) (40) encircling dsDNA. DNA binding residues are labeled, and aromatic wedge residues are indicated with black outlines. C. Crystal structure of the bovine papillomavirus E1 ATPase domain encircling ssDNA (PDB 2GXA) (8). D. Crystal structure of *Sulfolobus solfataricus* (SsoMCM) bound to ssDNA (PDB 6MII) (14).

Nevertheless, the engagement of Ile308 within the major groove is analogous to the aromatic wedge residue of H2i in MCM, raising the possibility that this aliphatic side chain acts as a steric wedge to partially destabilize the duplex. Whether this substitution supports equivalent unwinding efficiency, and what evolutionary pressures drove H2i loss and PS1 elongation in SaPI- and SCC-encoded helicases, remain open questions. Alternatively, H2i may have been added to MCM family helicases to compensate for a shorter PS1, or H2i and the elongated PS1 may have been independently added as they diverged from a common ancestor that lacked either of these features.

The duplex-bound single-hexamer intermediate captured here is structurally distinct from both the ssDNA-bound translocation intermediates reported for SaPI1 Rep and the duplex-encircling double hexamers observed in papillomavirus E1, SV40 T-antigen (LTag), and archaeal/eukaryotic MCMs (14, 17, 19, 20, 32, 33, 35, 39, 41, 42). The intermediate A-B form of the duplex near the base, transitioning to a more A-form character toward the top of the staircase, resembles the duplex DNA distortion imposed collectively at the interface of double hexamers, yet partially occurs here within a single hexamer. Whether this reflects early stage of DNA shearing mechanism by a pre-assembled single ring, a conformation favored by the short self-complementary substrate used in this study, or a transient intermediate preceding double hexamer assembly cannot be resolved from the current data. The preferential affinity of SaPIbov1 Rep for duplex DNA or ssDNA-dsDNA junctions suggested by these structures, combined with prior biochemical reports that origin loading and melting for SaPI-encoded Reps and MCMs can occur independently of ATP hydrolysis (49, 72, 73), raises the possibility that the intermediate captured here may reflect one of the intermediate pre-replicative states prior to double hexamer formation. How the loaded single hexamer transitions into an active unwinding configuration, and whether SaPIbov1 Rep assembles into double hexamers analogous to MCM, are important questions for future investigation.

A key mechanistic insight emerging from 3DVA is the PL1-PS1 conformational gate operating near the base of the staircase. Under non-hydrolyzing conditions, PL1 of the base subunit alternates between helical and horseshoe conformations in a manner structurally analogous to that reported for MCM, and these transitions are tightly coupled to nucleotide binding, intersubunit tightness, and the positional state of PS1 in the adjacent seam subunit. Specifically, helical PL1-DNA engagement permits downward insertion of the neighboring PS1 hairpin into the DNA groove in a geometry resembling the upper staircase subunits, where Ile308 is positioned for steric wedge activity, whereas horseshoe PL1 disengages from DNA and sterically precludes this insertion or displaces PS1 from the groove. Concurrently, PS1 ordering at the base of the staircase is reciprocally coupled to PS1 disorder at the top, reflecting coordinated protein-DNA interactions in response to positional shifts of DNA within the ring. The coexistence of a nucleotide occupancy gradient, asymmetric DNA engagement across subunits, and transient protein-protein and protein-DNA interactions observed within the data further supports that the helicase, captured under non-hydrolyzing conditions, does not represent a single artificially trapped conformation, but instead reflects a physiologically relevant ensemble of on-pathway translocation intermediates.

Based on these observations, we propose the following model for processive translocation (Figure 7). In the initiating state, PS1 of the topmost staircase subunit is partially engaged with the incoming DNA substrate while the horseshoe PL1 at the base is partially disengaged. Helical engagement of PL1 with DNA at the base permits repositioning of the PS1 of the adjacent seam subunit into the groove, cooperatively capturing DNA in an advanced register. This capture is coupled to PS1 disengagement at the top of the staircase, promoting ATP binding at the newly tightened upper subunit interface and ADP release at the relaxed lower subunit interface. Nucleotide exchange drives intersubunit reorganization, advancing the seam by one staircase step and resetting the PL1-PS1 gate for the next translocation step. Iterative progression of this cycle, coupled with ATP hydrolysis, constitutes a hand-over-hand mechanism for processive dsDNA translocation with duplex unwinding mediated by the steric wedge. Direct validation of this model will require structural characterization of duplex bound single or double hexamer assemblies under ATP hydrolyzing conditions.

**Figure 7.**
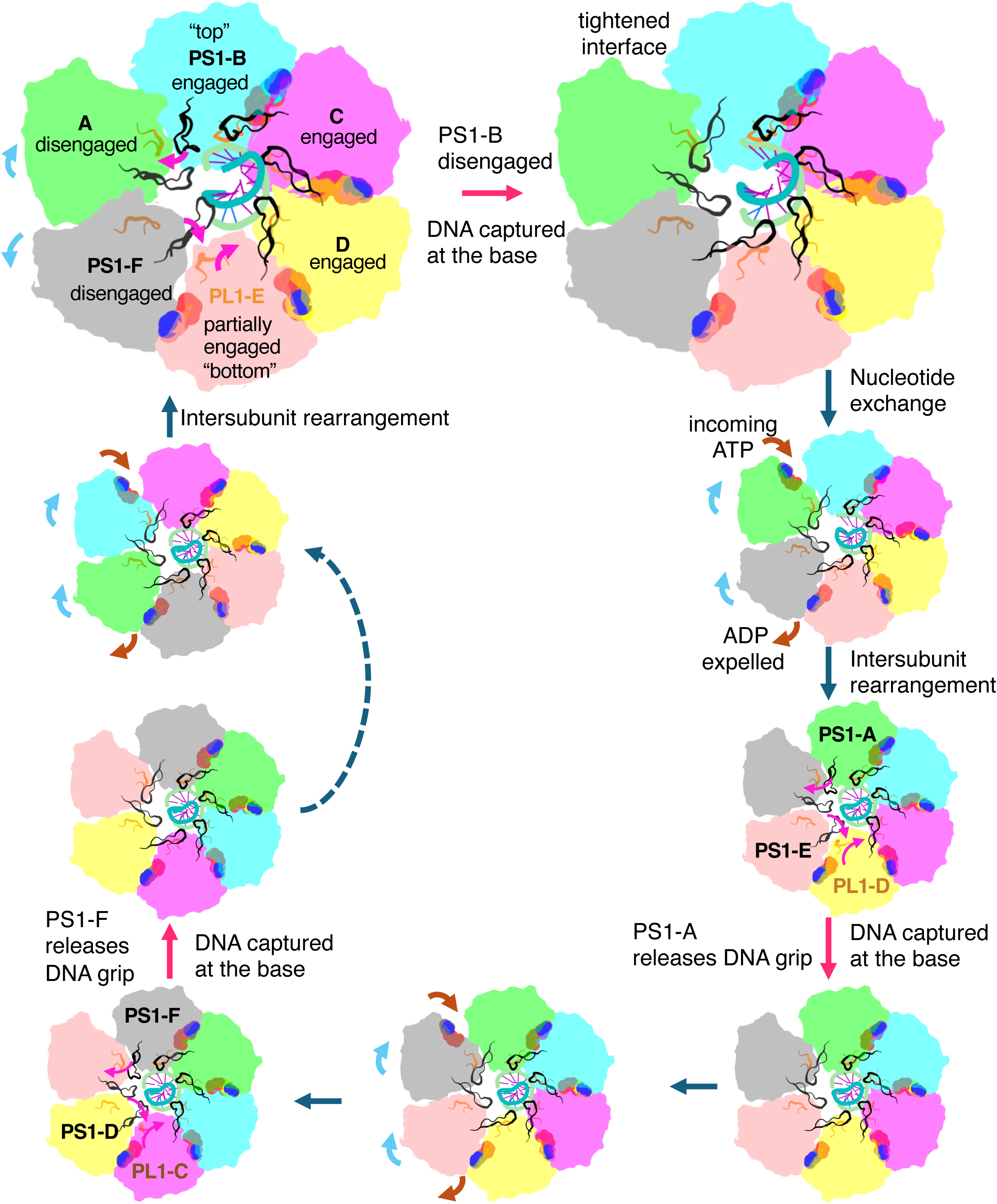
Schematic model of ATP-coupled DNA translocation by SaPIbov1 Rep. A schematic pathway for SaPIbov1 Rep DNA translocation, with protein, ATP, and DNAs colored as in previous figures. The pathway begins with Rep encircling dsDNA in spiral staircase arrangement, with subunits assigned to the top, middle, and bottom positions. ATP is bound at the top to middle interfaces, whereas ADP is bound at the bottom of the staircase. As DNA enters the ring, PS1-B at the top engages the incoming leading strand and extends stabilization along the staircase. Seam subunits (A and F) exhibit transient interactions with the neighboring subunits or DNA. Upon opening of the seam interface (indicated with blue arrows), DNA gains conformational freedom. PL1-E engages the shifted DNA, allowing PS1-F to complete DNA capture as PS1-B at the top disengages. PS1-B is then transiently stabilized against the PL1-A, forming a tightened interface for ATP binding at subunit A. ATP binding at the interface A-B drives intersubunit rearragnment, while the ADP bound E-F interface destabilizes, likely releasing ADP and advancing the ATPase cycle with DNA translocation.

Comparison with other SaPI-encoded Reps highlights divergent mechanistic strategies. Cryo-EM structures of SaPI1 and SaPI5 Reps, which harbor SF3-family ATPase domains, exhibit pronounced CTD flexibility proposed to be essential for ATPase activity and productive helicase function. SaPIbov1 Rep, by contrast, displays a comparatively immobile CTD, with conformational plasticity instead concentrated at the seam subunits of the ATPase domain. This redistribution of flexibility may reflect an intrinsic difference between SF3 and SF6 ATPase architectures in how conformational dynamics are partitioned across the ring, or may alternatively indicate that SaPIbov1 Rep was captured in a duplex preferred state in which CTD engagement with substrate has not yet occurred. Notably, Pri-Rep complex formation, mediated through the N-terminal helix of Rep, was required to overcome severe orientation bias during cryo-EM data collection, yet Pri density was absent in the final reconstruction, suggesting that the primase dissociated during grid preparation or was disordered. The absence of Pri from the final maps precludes conclusions about its structural role in modulating Rep conformation. However, the conservation of this interaction interface in SaPI2 Rep, but not in Cch, suggests that SCC and SaPI elements have evolved distinct strategies for priming DNA replication. Resolving whether the duplex threading mode captured here is a functionally productive intermediate or a kinetically trapped state will require reconstitution with *ori* DNA and under conditions permitting ATP hydrolysis.

Further questions include how SaPIbov1 Rep transitions from recognizing the 12 individual iterons within its cognate ori to two hexamers encircling DNA (Supplementary Figure 7A). Within ori, many pairs or trios of iterons are approximately one helical turn of DNA apart (Supplementary Figure 7B). This arrangement suggests that binding of the C-terminal WH domains to the first 3 iterons would place the beginning of those domains 35-38 Å apart – remarkably close to their 33-34 Å spacing in the assembled hexamer (Supplementary Figure 7B and D). Ori binding by individual subunits’ C-terminal domains may therefore facilitate formation of the ATPase and N-terminal domain protein-protein interfaces, perhaps concomitantly with dissociation of the C-terminal domains from the origin DNA and rotation about the flexible linker that connects them to the ATPase to assemble the C-terminal domain ring seen in the hexamer. The crystal structure of the structurally-related TcpK bound to repeated iterons showed that TcpK dimerization brings two DNAs into close proximity (Supplementary Figure 7F). AlphaFold3 predicted of Rep C-terminal WH domain dimers inconsistently, but if such dimers did form, they could bring together distal regions of the origin (perhaps with the help of a DNA bending protein), thus bringing all required Rep subunits into close proximity (Supplementary Figure 7C).

Collectively, these structures establish a framework for understanding how a minimal self-loading SaPI replication initiator engages and distorts duplex DNA after hexamer formation but prior to replication. The elongated PS1 motif, the PL1-PS1 conformational gate, and the dynamic seam subunits together define an alternative structural solution for helicase function that diverges from the canonical MCM mechanism while preserving its core structural elements for nucleotide-coupled DNA binding and translocation. These findings motivate future biochemical and structural studies to resolve how SaPIbov1 Rep loads onto the origin, whether it forms double hexamers, and how it can initiate the transition from a duplex-bound pre-replicative state to active strand separation.

## Data Availability

- All supplemental movies have been deposited in the Mendeley Data repository at the following link: (https://doi.org/10.17632/tkzpwdgnk4.1).
- The x-ray diffraction data has been deposited at SBgrid: data sets #1033 (native) and #1034 (SeMet).
- The raw cryo-EM images have been deposited at the Electron Microscopy Public Image Archive (EMPIAR) under code EMPIAR-13092.
- The models generated in this study have been deposited at the Protein Data Bank (PDB) with the following accession codes PDB: 8TCH (apo structure), 39QS, 39QT, 39QU, 39QW, 39QZ, 39QX, and 39RB.
- Cryo-EM maps have been deposited in the Electron Microscopy Data Bank (EMDB) under codes EMDB: EMD-79702, EMD-79703, EMD-79705, EMD-79708, EMD-79710, EMD-79709, and EMD-79712.

## Supplementary Data

## Author Contributions

Conception and design, Methodology: PAR, HS, IM-S

Funding acquisition: PAR

Data acquisition: HS, IM-S, YP

Data analysis: PAR, HS

Drafting and revision: PAR, HS

## Acknowledgements

We are grateful for access to the GM/CA 23-ID-B and 23-ID-D beamlines staff for support in X-ray data collection. GM/CA at Advanced Photon Source has been funded by the National Cancer Institute (ACB-12002) and the National Institute of General Medical Sciences (AGM-12006, P30GM138396). This research used resources of the Advanced Photon Source, a U.S. Department of Energy (DOE) Office of Science User Facility operated for the DOE office of Science by Argonne National Laboratory under Contract No. DE-AC02-06CH11357. The Eiger 16M detector was funded by NIH grant S10OD012289. We thank the members of Research Computing Center (RCC) at the University of Chicago for providing their services. This work was completed using the Beagle3 high-performance computing cluster funded by the National Institutes of Health through grant 1S10OD028655-01. We thank James R. Fuller for his assistance in cryo-EM data acquisition; Cryo-EM datasets were collected at the University of Chicago Advanced Electron Microscopy Core Facility (RRID:SCR_019198). Molecular graphics and analyses performed with UCSF ChimeraX, developed by the Resource for Biocomputing, Visualization, and Informatics at the University of California, San Francisco, with support from National Institutes of Health R01-GM129325 and the Office of Cyber Infrastructure and Computational Biology, National Institute of Allergy and Infectious Diseases; compiled and configured by SBGrid (74).

---

**Extended Data Table 1.**
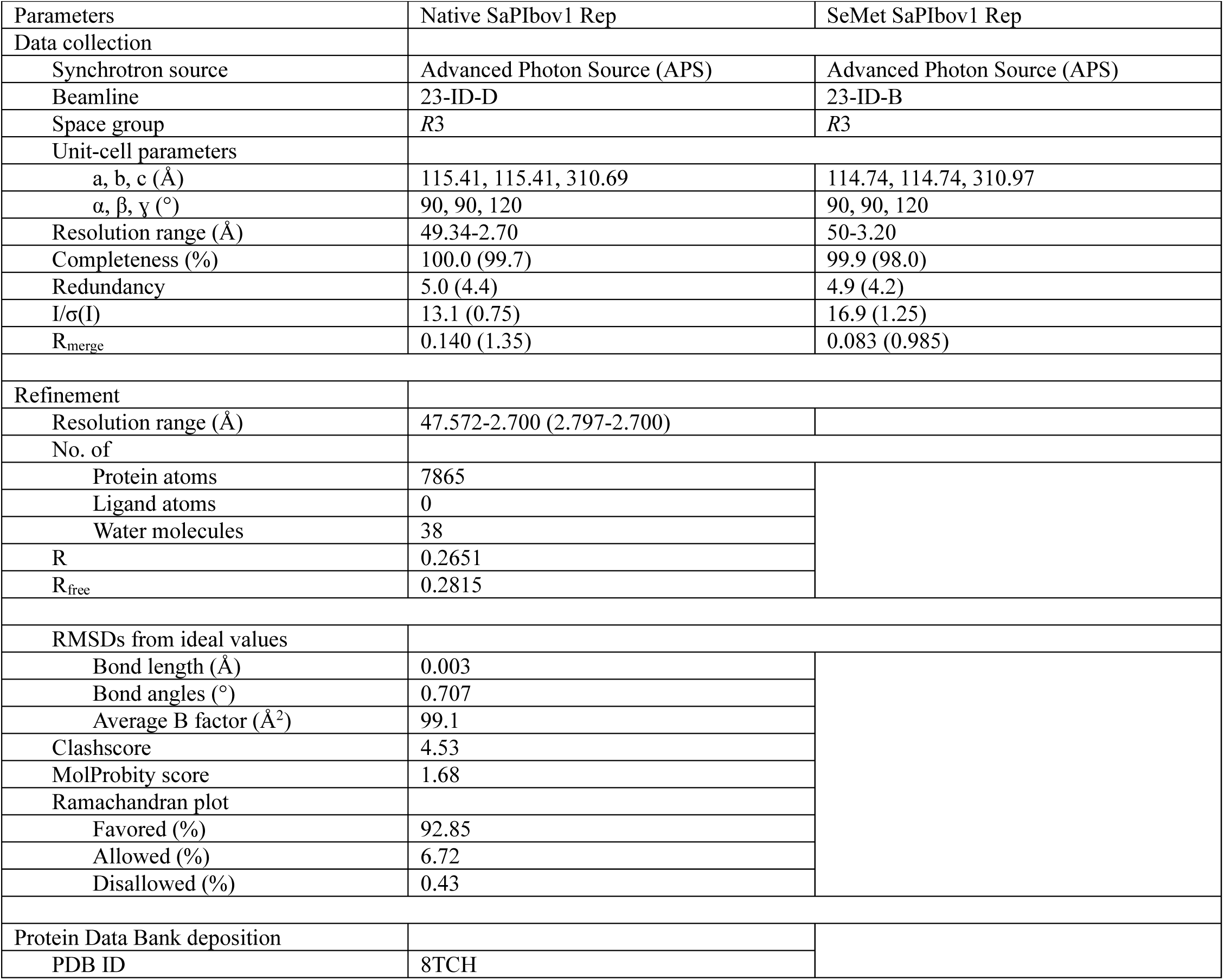
X-ray diffraction data collection and refinement statistics.

| Parameters | Native SaPIbov1 Rep | SeMet SaPIbov1 Rep |
|---|---|---|
| Data collection |  |  |
| Synchrotron source | Advanced Photon Source (APS) | Advanced Photon Source (APS) |
| Beamline | 23-ID-D | 23-ID-B |
| Space group | *R*3 | *R*3 |
| Unit-cell parameters |  |  |
| a, b, c (Å) | 115.41, 115.41, 310.69 | 114.74, 114.74, 310.97 |
| α, β, ɣ (°) | 90, 90, 120 | 90, 90, 120 |
| Resolution range (Å) | 49.34-2.70 | 50-3.20 |
| Completeness (%) | 100.0 (99.7) | 99.9 (98.0) |
| Redundancy | 5.0 (4.4) | 4.9 (4.2) |
| I/σ(I) | 13.1 (0.75) | 16.9 (1.25) |
| R<sub>merge</sub> | 0.140 (1.35) | 0.083 (0.985) |
| Refinement |  |  |
| Resolution range (Å) | 47.572-2.700 (2.797-2.700) |  |
| No. of |  |  |
| Protein atoms | 7865 |  |
| Ligand atoms | 0 |  |
| Water molecules | 38 |  |
| R | 0.2651 |  |
| R<sub>free</sub> | 0.2815 |  |
| RMSDs from ideal values |  |  |
| Bond length (Å) | 0.003 |  |
| Bond angles (°) | 0.707 |  |
| Average B factor (Å<sup>2</sup>) | 99.1 |  |
| Clashscore | 4.53 |  |
| MolProbity score | 1.68 |  |
| Ramachandran plot |  |  |
| Favored (%) | 92.85 |  |
| Allowed (%) | 6.72 |  |
| Disallowed (%) | 0.43 |  |
| Protein Data Bank deposition |  |  |
| PDB ID | 8TCH |  |

**Extended Data Table 2.**
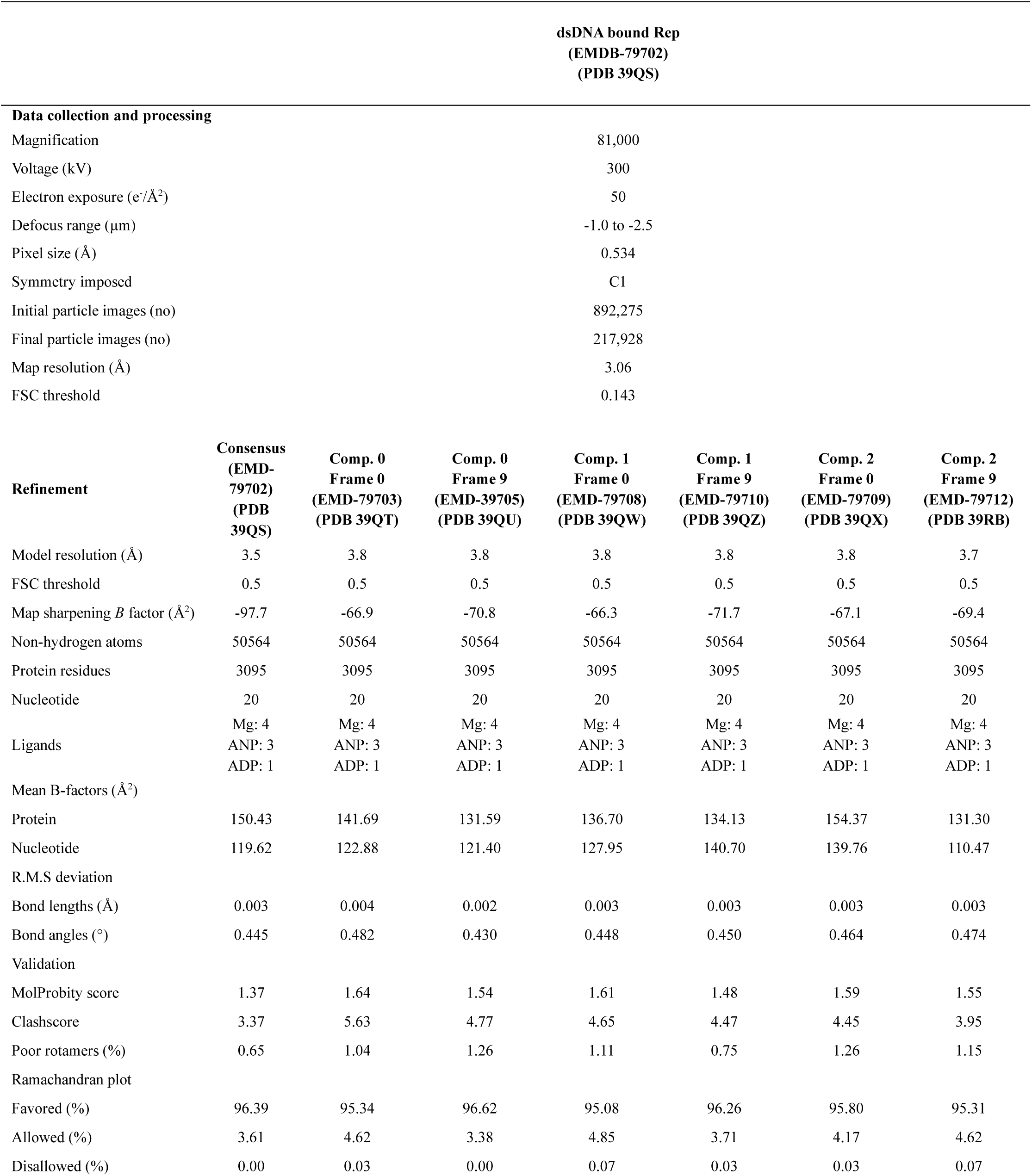
Cryo-EM Data Collection, Refinement and Validation Statistics.

|  | dsDNA bound Rep<br>(EMDB-79702)<br>(PDB 39QS) |  |  |  |  |  |  |
|---|---|---|---|---|---|---|---|
| **Data collection and processing** |  |  |  |  |  |  |  |
| Magnification | 81,000 |  |  |  |  |  |  |
| Voltage (kV) | 300 |  |  |  |  |  |  |
| Electron exposure (e<sup>-</sup>/Å<sup>2</sup>) | 50 |  |  |  |  |  |  |
| Defocus range (µm) | -1.0 to -2.5 |  |  |  |  |  |  |
| Pixel size (Å) | 0.534 |  |  |  |  |  |  |
| Symmetry imposed | C1 |  |  |  |  |  |  |
| Initial particle images (no) | 892,275 |  |  |  |  |  |  |
| Final particle images (no) | 217,928 |  |  |  |  |  |  |
| Map resolution (Å) | 3.06 |  |  |  |  |  |  |
| FSC threshold | 0.143 |  |  |  |  |  |  |
| **Refinement** | **Consensus<br>(EMD-79702)<br>(PDB 39QS)** | **Comp. 0<br>Frame 0<br>(EMD-79703)<br>(PDB 39QT)** | **Comp. 0<br>Frame 9<br>(EMD-39705)<br>(PDB 39QU)** | **Comp. 1<br>Frame 0<br>(EMD-79708)<br>(PDB 39QW)** | **Comp. 1<br>Frame 9<br>(EMD-79710)<br>(PDB 39QZ)** | **Comp. 2<br>Frame 0<br>(EMD-79709)<br>(PDB 39QX)** | **Comp. 2<br>Frame 9<br>(EMD-79712)<br>(PDB 39RB)** |
| Model resolution (Å) | 3.5 | 3.8 | 3.8 | 3.8 | 3.8 | 3.8 | 3.7 |
| FSC threshold | 0.5 | 0.5 | 0.5 | 0.5 | 0.5 | 0.5 | 0.5 |
| Map sharpening *B* factor (Å<sup>2</sup>) | -97.7 | -66.9 | -70.8 | -66.3 | -71.7 | -67.1 | -69.4 |
| Non-hydrogen atoms | 50564 | 50564 | 50564 | 50564 | 50564 | 50564 | 50564 |
| Protein residues | 3095 | 3095 | 3095 | 3095 | 3095 | 3095 | 3095 |
| Nucleotide | 20 | 20 | 20 | 20 | 20 | 20 | 20 |
| Ligands | Mg: 4<br>ANP: 3<br>ADP: 1 | Mg: 4<br>ANP: 3<br>ADP: 1 | Mg: 4<br>ANP: 3<br>ADP: 1 | Mg: 4<br>ANP: 3<br>ADP: 1 | Mg: 4<br>ANP: 3<br>ADP: 1 | Mg: 4<br>ANP: 3<br>ADP: 1 | Mg: 4<br>ANP: 3<br>ADP: 1 |
| Mean B-factors (Å<sup>2</sup>) |  |  |  |  |  |  |  |
| Protein | 150.43 | 141.69 | 131.59 | 136.70 | 134.13 | 154.37 | 131.30 |
| Nucleotide | 119.62 | 122.88 | 121.40 | 127.95 | 140.70 | 139.76 | 110.47 |
| R.M.S deviation |  |  |  |  |  |  |  |
| Bond lengths (Å) | 0.003 | 0.004 | 0.002 | 0.003 | 0.003 | 0.003 | 0.003 |
| Bond angles (°) | 0.445 | 0.482 | 0.430 | 0.448 | 0.450 | 0.464 | 0.474 |
| Validation |  |  |  |  |  |  |  |
| MolProbity score | 1.37 | 1.64 | 1.54 | 1.61 | 1.48 | 1.59 | 1.55 |
| Clashscore | 3.37 | 5.63 | 4.77 | 4.65 | 4.47 | 4.45 | 3.95 |
| Poor rotamers (%) | 0.65 | 1.04 | 1.26 | 1.11 | 0.75 | 1.26 | 1.15 |
| Ramachandran plot |  |  |  |  |  |  |  |
| Favored (%) | 96.39 | 95.34 | 96.62 | 95.08 | 96.26 | 95.80 | 95.31 |
| Allowed (%) | 3.61 | 4.62 | 3.38 | 4.85 | 3.71 | 4.17 | 4.62 |
| Disallowed (%) | 0.00 | 0.03 | 0.00 | 0.07 | 0.03 | 0.03 | 0.07 |

## Supplementary figures

**Supplementary figure 1.**
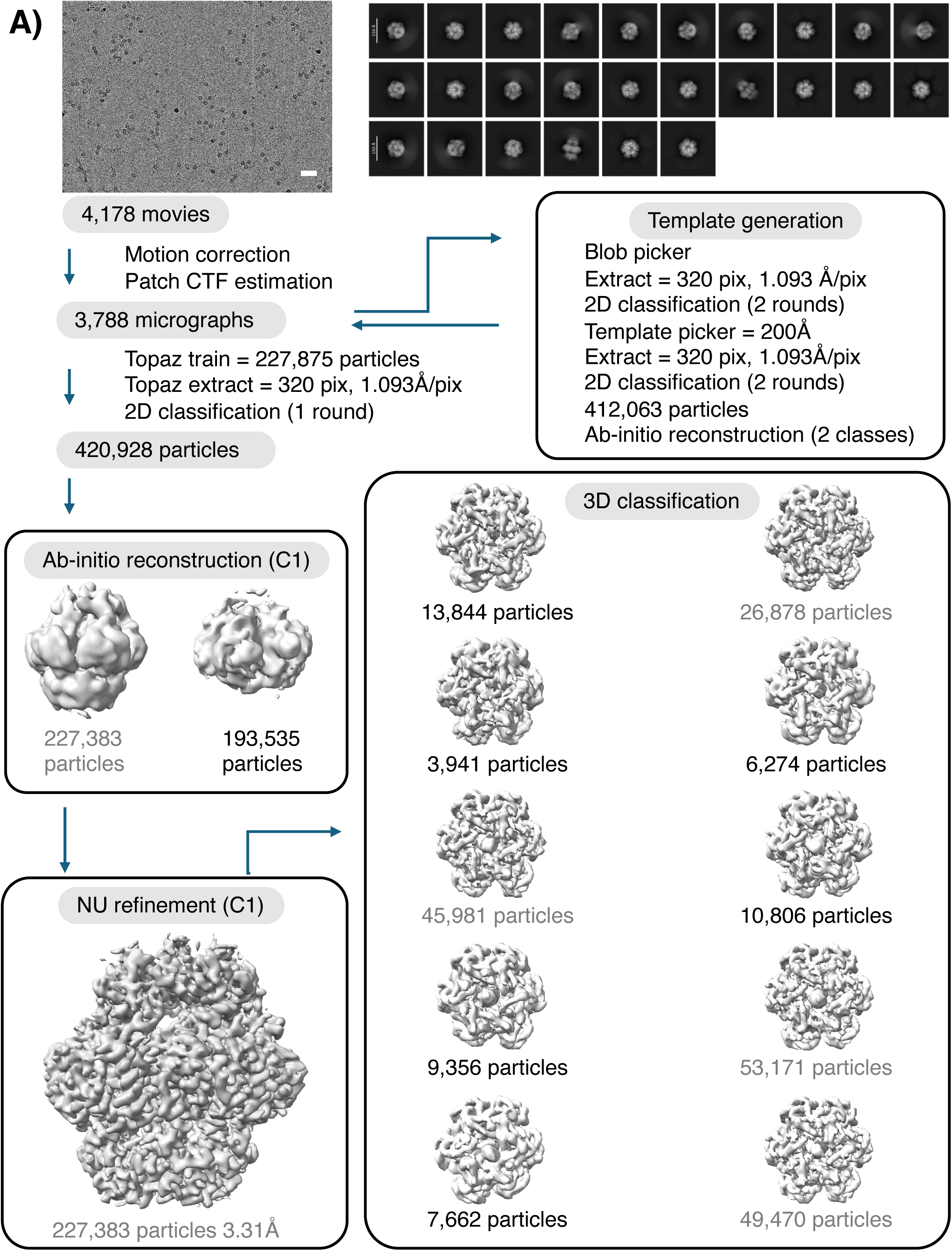
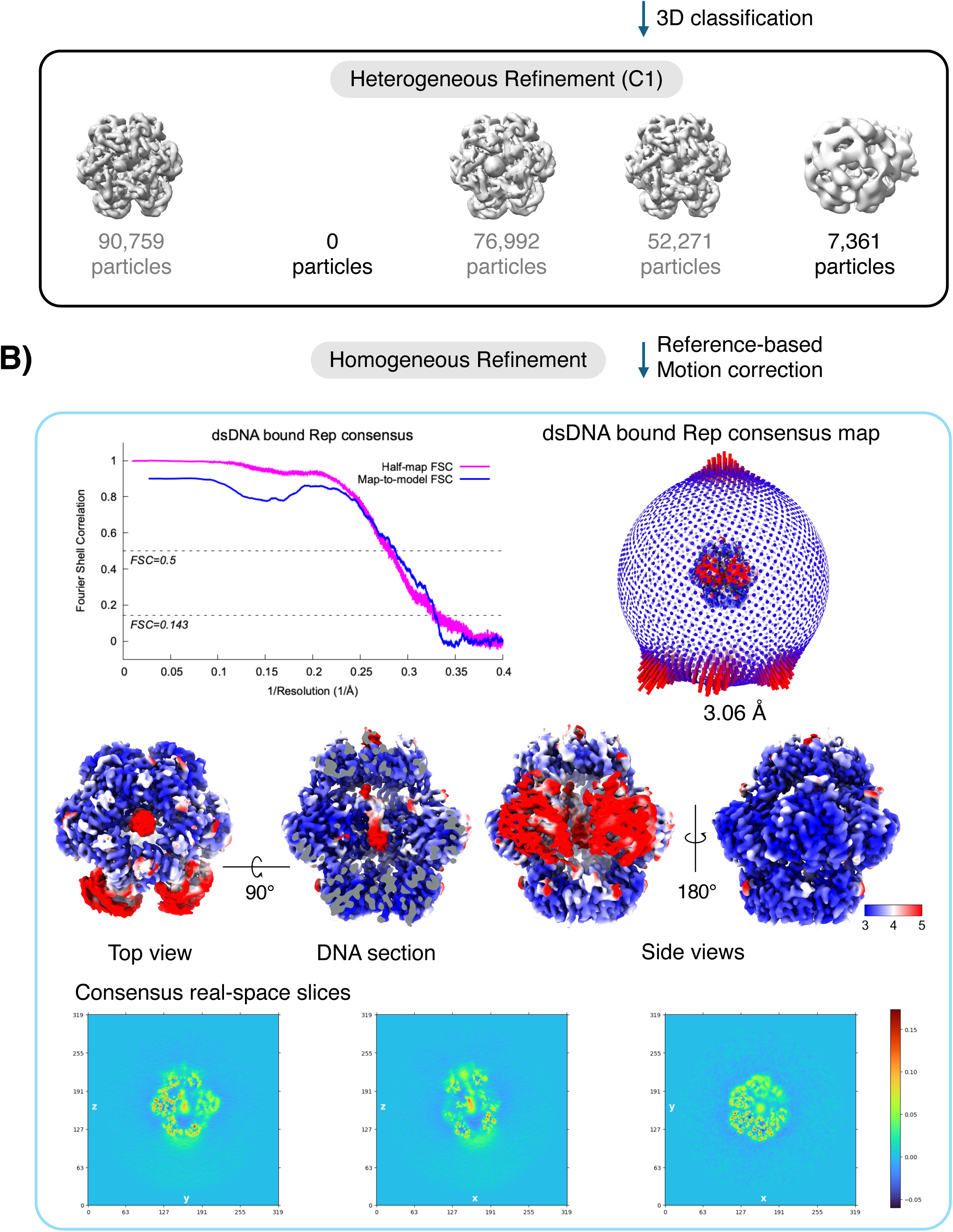
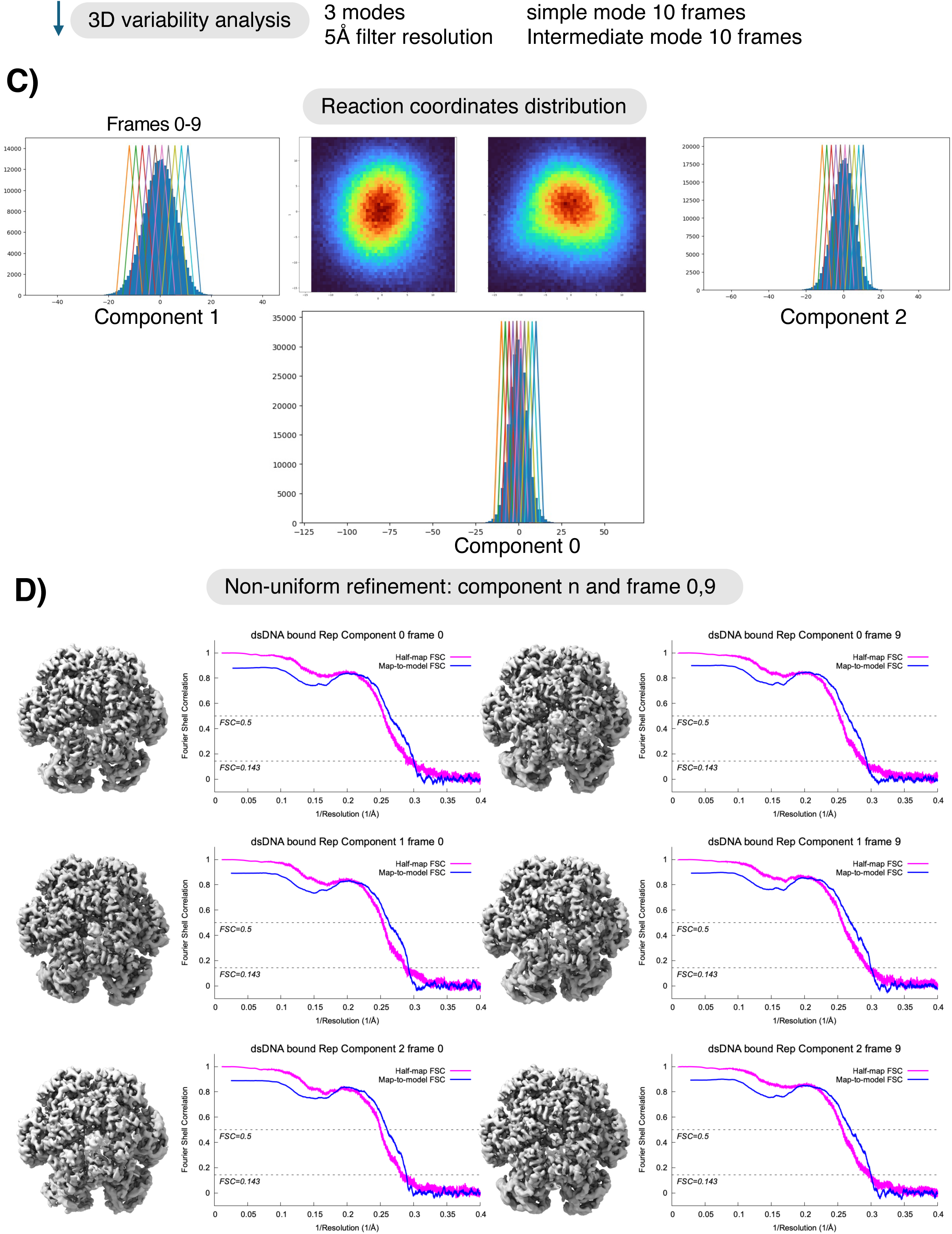
Data processing flow chart for the dsDNA bound SaPIbov1 Rep data. A. A visual overview of the data processing steps of dsDNA bound SaPIbov1 Rep dataset. The dataset was subjected to particle selection, iterative rounds of 2D classification, 3D classification and heterogeneous refinement. A representative micrograph (scale bar, 45nm) and representative 2D class averages are shown. B. The distribution of the Euler angles is shown next to the consensus map generated from homogeneous refinement. Four views of the consensus map are colored by local resolution. Fourier shell correlation (FSC) curve of the masked map after cryoSPARC (61) postprocessing. The resolution was determined by the FSC=0.143 (magenta). The map-to-model FSC curve is also shown (blue). C. 3D variability analysis (3DVA) (56) was performed using cryoSPARC (61) with three modes and 5Å resolution filter configuration. 20 linear traversal volumes per principal component (PC) were generated for movies using ‘simple mode’ in 3DVA display. 10 volumes per PC were generated for further modeling using ‘intermediate mode’ in 3DVA display. D. The first and last frames of each PC were used to generate the final maps using non-uniform refinement. The fourier shell correlation (FSC) curve of the masked map (magenta) and the map-to-model FSC curve (blue) are shown next to the maps.

**Supplementary figure 2.**
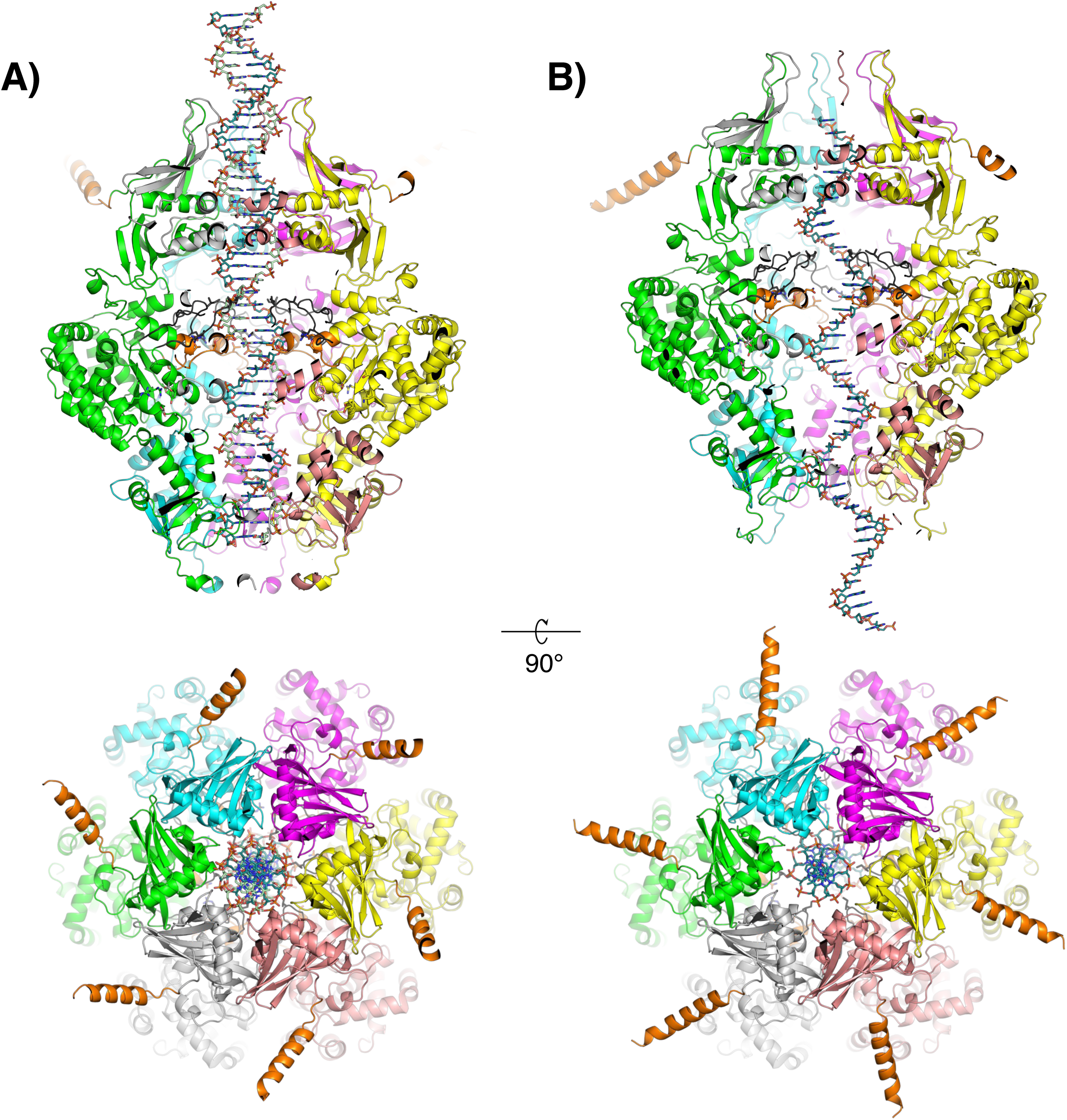
AlphaFold3-predicted model of SaPIbov1 Rep hexamer with ssDNA and dsDNA. A. AlphaFold3 predicted structure of SaPIbov1 Rep with dsDNA threaded through the central channel, shown in sliced side (top) and top (bottom) views. B. Similar views of SaPIbov1 Rep with ssDNA.

**Supplementary figure 3.**
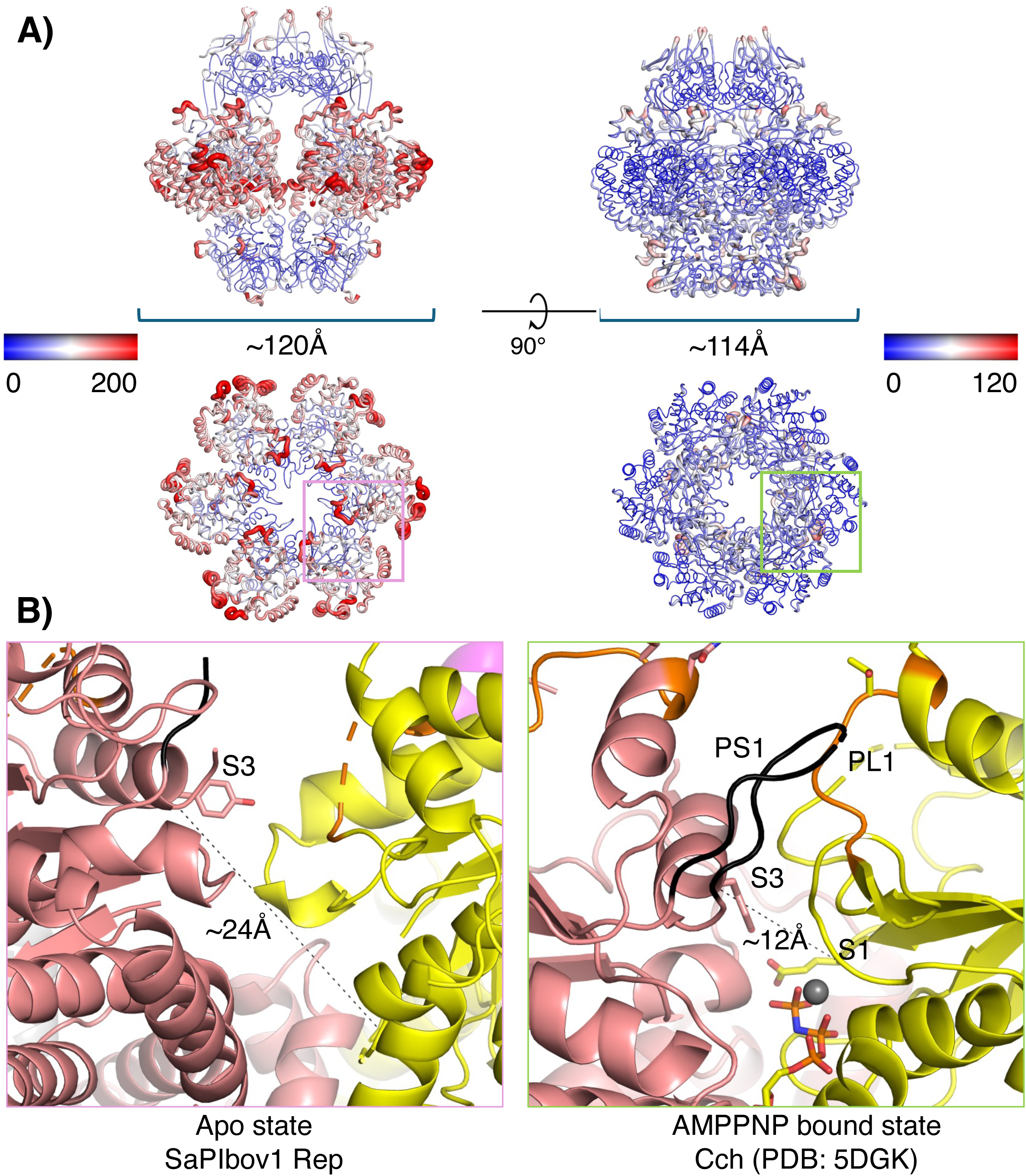
Structural comparison of SaPIbov1 Rep with SCC*mec* Cch. A. Structures of SaPIbov1 Rep (left panels) and SCC*mec* Cch (53) (right panels, PDB 5DGK) shown in side (top) and top (bottom) views, colored by B-factors (ranges: 0 to 200 for Rep and 0 to 120 for Cch). B. Close-up views of ATPase nucleotide-binding sites: an empty binding site in SaPIbov1 Rep (pink box in panel A) verses an AMPPNP bound site in SCC*mec* Cch (53) (green box in panel A). The distance between Cα atoms of S3 and S1 is displayed by a black dashed line.

**Supplementary figure 4.**
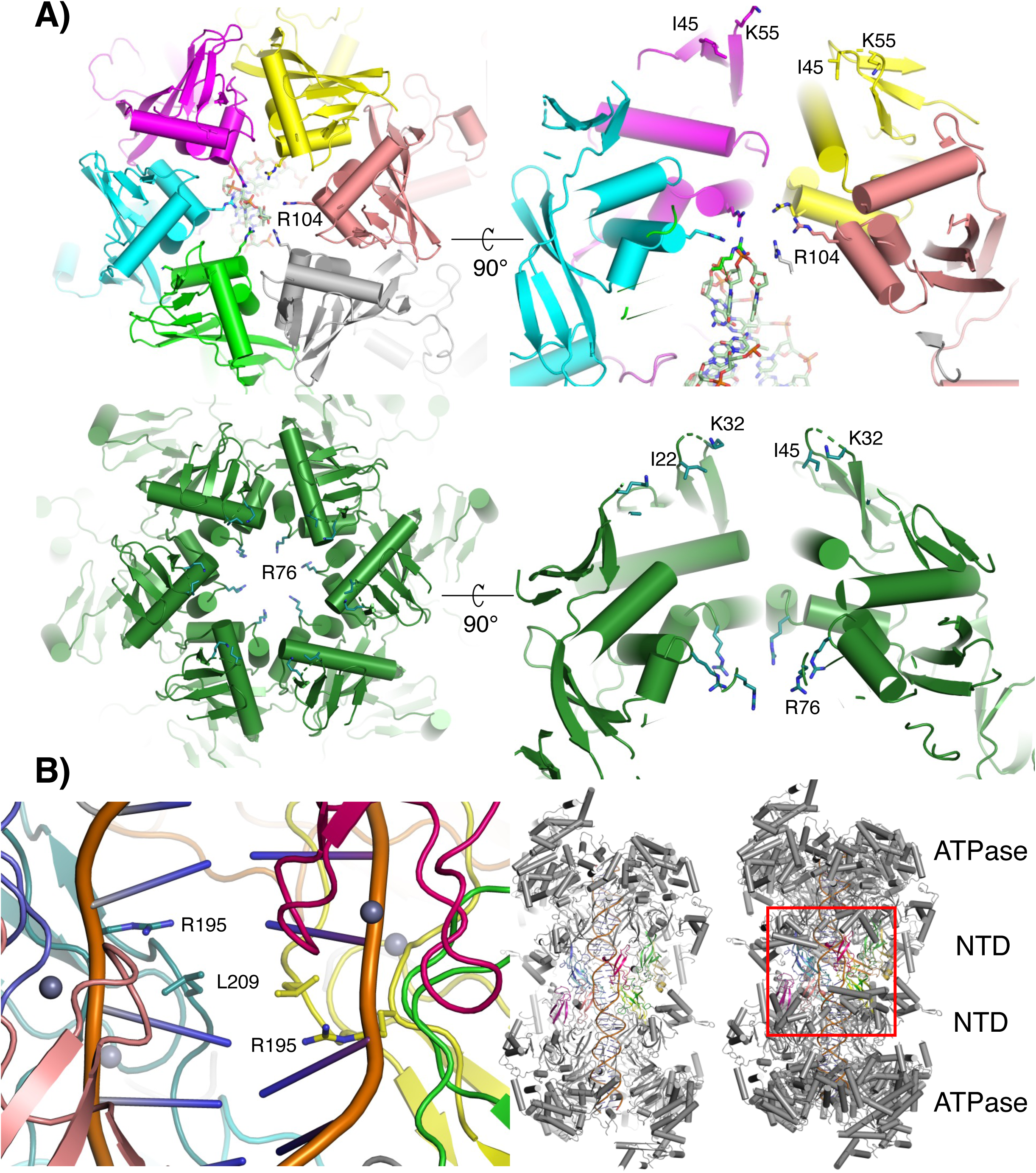
Structural comparison of SaPIbov1 Rep with SCC*mec* Cch and MCM NTDs. A. Close-up views of N-terminal domains (NTD): SaPIbov1 Rep (top row; left top view; right, sliced side view) and Cch (PDB 5DGK) (53) green; bottom row; left, top view; right, sliced side view). Residues lining the NTD channel are shown as sticks. B. Close-up view of the MCM double hexamer (PDB 8S09) (41) interface (boxed region in right panel), highlighting intercalated residues R195 and L209 from MCM5.

**Supplementary figure 5.**
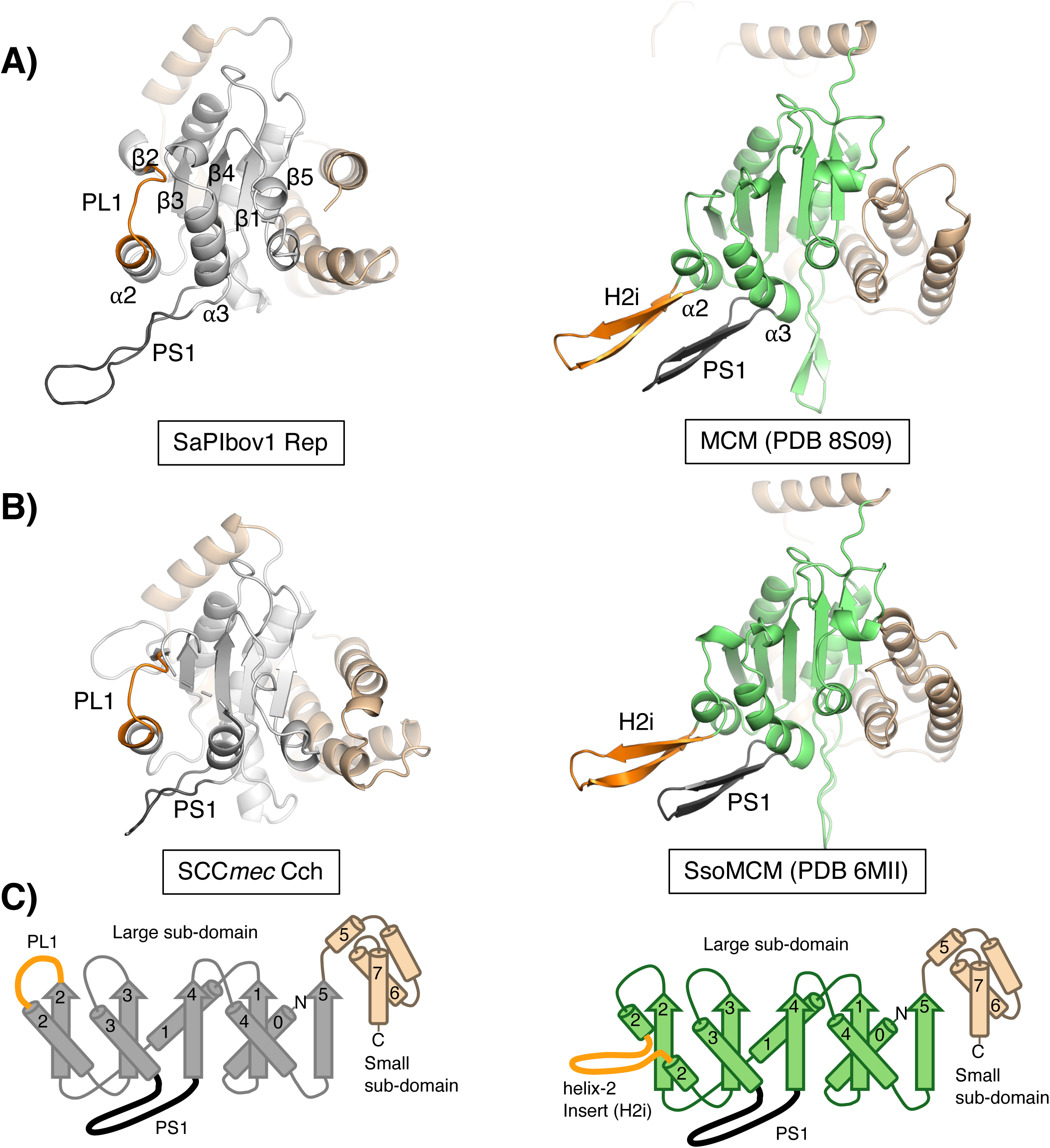
Structural comparison of SaPIbov1 Rep and MCM ATPase domains. A. A close-up views of ATPase domains: SaPIbov1 Rep (left) and MCM (right; PDB 8S09) (41). Secondary structure elements are labeled as in panel C. PL1 and H2i motifs are shown in orange, and PS1 motifs in black. B. Cch (left; PDB 5DGK) (53) and SsoMCM (right; PDB 6MII) (14). Same orientation and color scheme as panel C. C. Topology diagrams of ATPase domains from SaPIbov1 Rep (left) and MCM-like superfamily (right).

**Supplementary figure 6.**
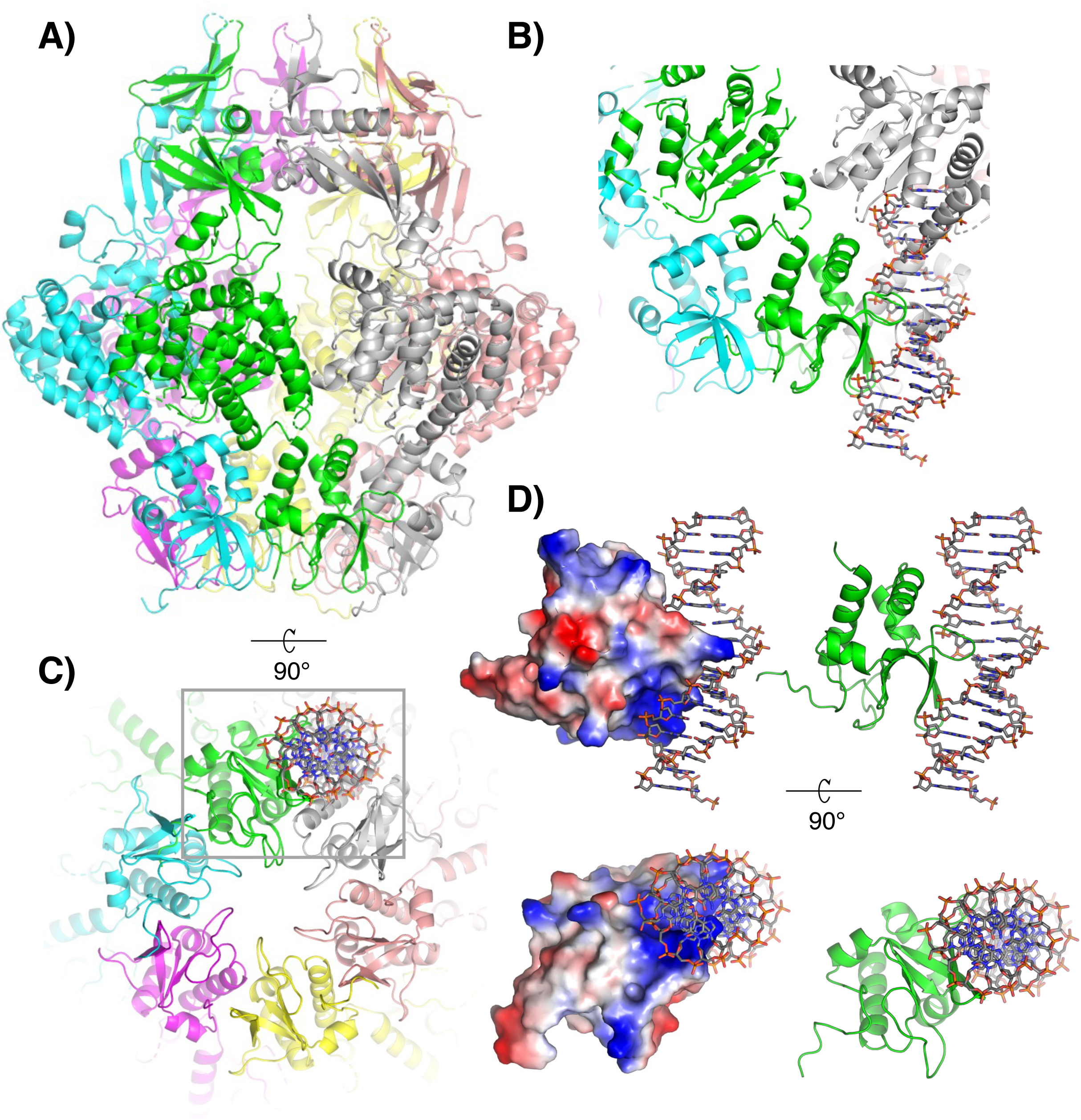
AlphaFold3-predicted SaPIbov1 Rep CTD with dsDNA. A. Structure of SaPIbov1 Rep shown in side view. B. Close-up view of SaPIbov1 Rep superimposed with an AlphaFold3 (65) predicted CTD monomer bound to dsDNA. C. Top view of the superimposed structures (in panel B), showing the buried winged-helix (WH) dsDNA binding motif within the hexameric ring. D. AlphaFold3 (65) predicted CTD monomer with dsDNA shown as an electrostatic potential surface map in left panel (positively charged surface in blue, net 0 charge surface in white, and negatively charge surface in red).

**Supplementary figure 7.**
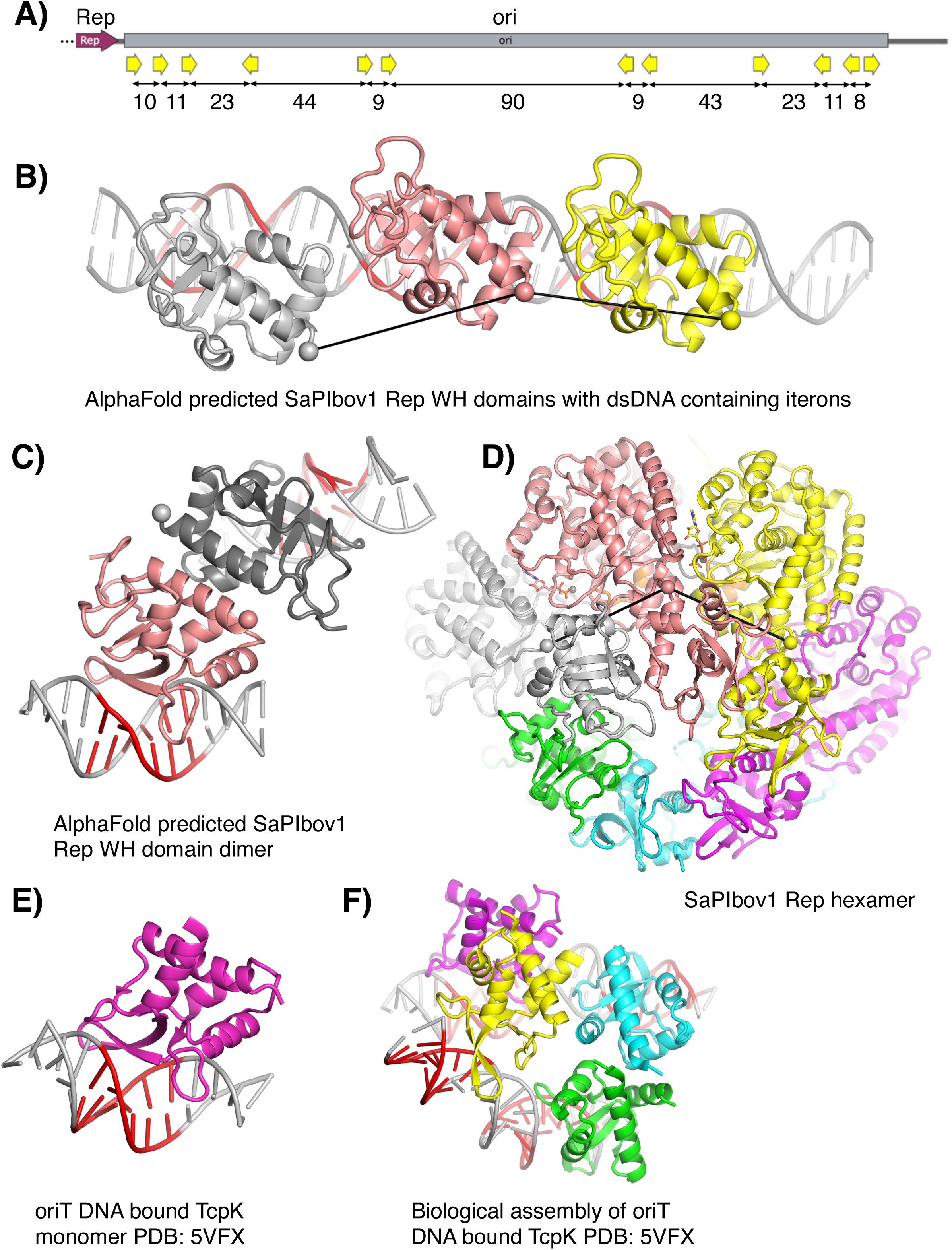
Comparison of the C-terminal WH domain of SaPIbov1 Rep with the pCW3 relaxosome protein TcpK. A. Diagram of the SaPIbov1 origin of replication, which immediately follows the Rep coding region. Iterations of the sequence GTACCC (or GTCCCC) are highlighted as yellow arrows, and the number of base pairs between their centers is noted below. B. AlphaFold3 (65) model of 3 copies of the SaPIbov1 Rep C-terminal WH domain bound to the first 3 iterons (perhaps due to homology to TcpK (69) which binds GTTCCC, AlphaFold3 did dock the proteins onto their cognate binding sites. Spheres mark the C*a* of residue 453 at the beginning of the C-terminal WH domain. These are 38 and 35Å apart, respectively. The corresponding iterons are colored red. C. AlphaFold3 (65) model of 2 copies of the SaPIbov1 Rep C-terminal WH domain bound to the two separate dsDNA containing iterons bringing two copies in close proximity. D. Experimental structure of the SaPIbov1 Rep hexamer, viewed from the C-terminal end. The DNA binding surface of C-terinal domains is occluded, but the spheres marking residue 453 are 33-34Å apart - very similar to the spacing in panel B. E. Close-up view of the pCW3 relaxosome protein TcpK monomer in the same orientation as in panel A. F. Biological assembly of the pCW3 relaxosome protein TcpK (PDB 5VFX) (69) bound to origin of transfer (oriT) DNA. The core region of tandem repeats (GTTCCC or GTTCCT) is colored in red.

**Supplementary figure 8.**
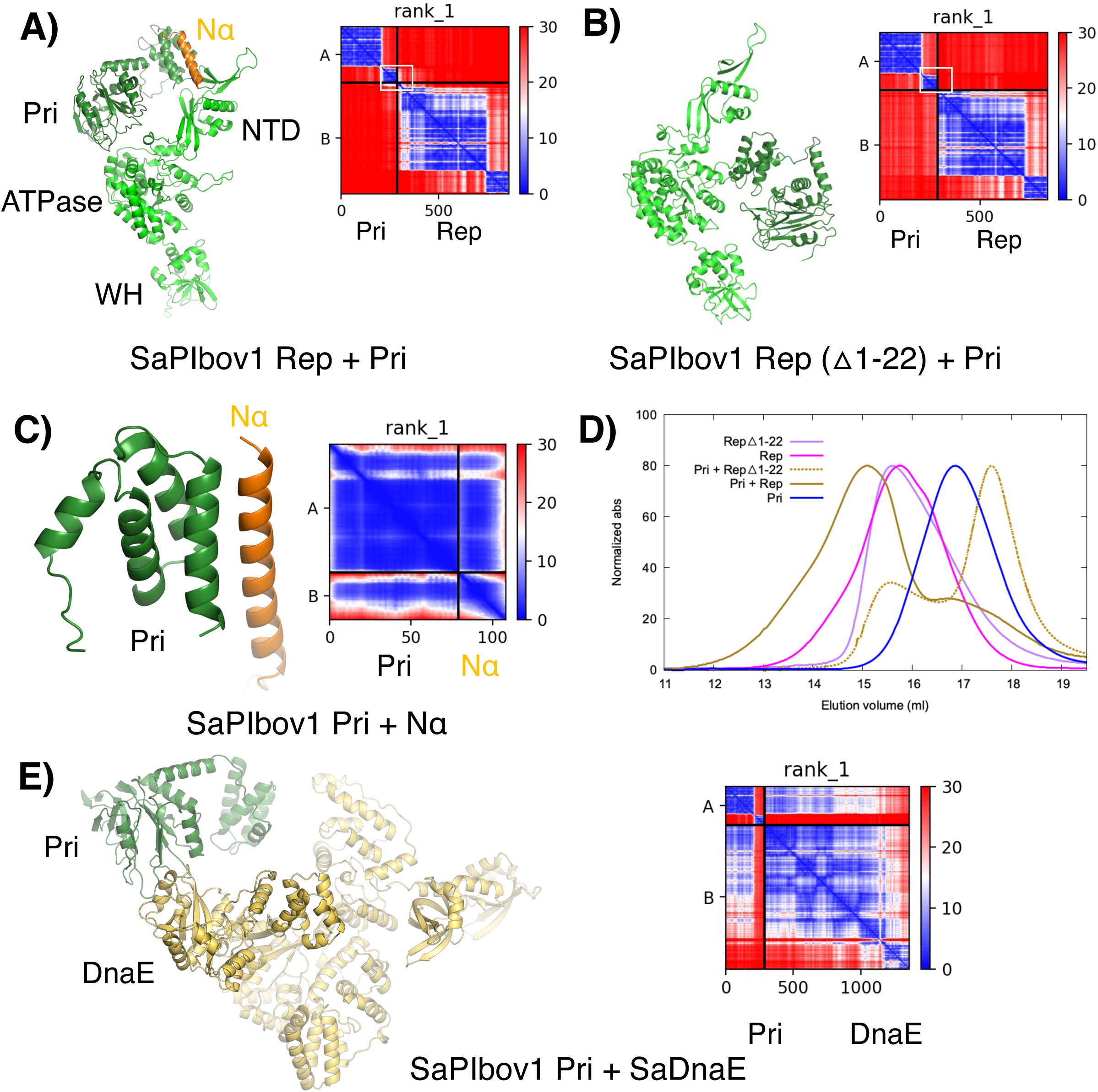
Nα of the NTD in SaPIbov1 Rep interacts with SaPIbov1 Pri: WT vs. △Nα (residues 1-22) A. AlphaFold2-multimer prediction of SaPIbov1 Rep WT(pale green) in complex with SaPIbov1 Pri (dark green) and the predicted aligned error (PAE) plot (first rank) shown on the right. Lower PAE values (blue) indicate higher confidence. B. AlphaFold2-multimer prediction of the △Nα (residues 1-22) mutant in complex with SaPIbov1 Pri, with corresponding PAE plot (first rank). C. AlphaFold2-multimer prediction of the C-terminal Pri (dark green) in complex with SaPIbov1 Rep Nα (orange, residues 1-22) with corresponding PAE plot (first rank). D. Size-exclusion chromatography (SEC) elution profiles of SaPIbov1 Rep WT and △Nα mutant in the presence and absence of SaPIbov1 Pri. E. AlphaFold2-multimer prediction of the SaPIbov1 Pri (dark green) and SaDnaE (yellow) with corresponding PAE plot (first rank).

**Supplementary figure 9.**
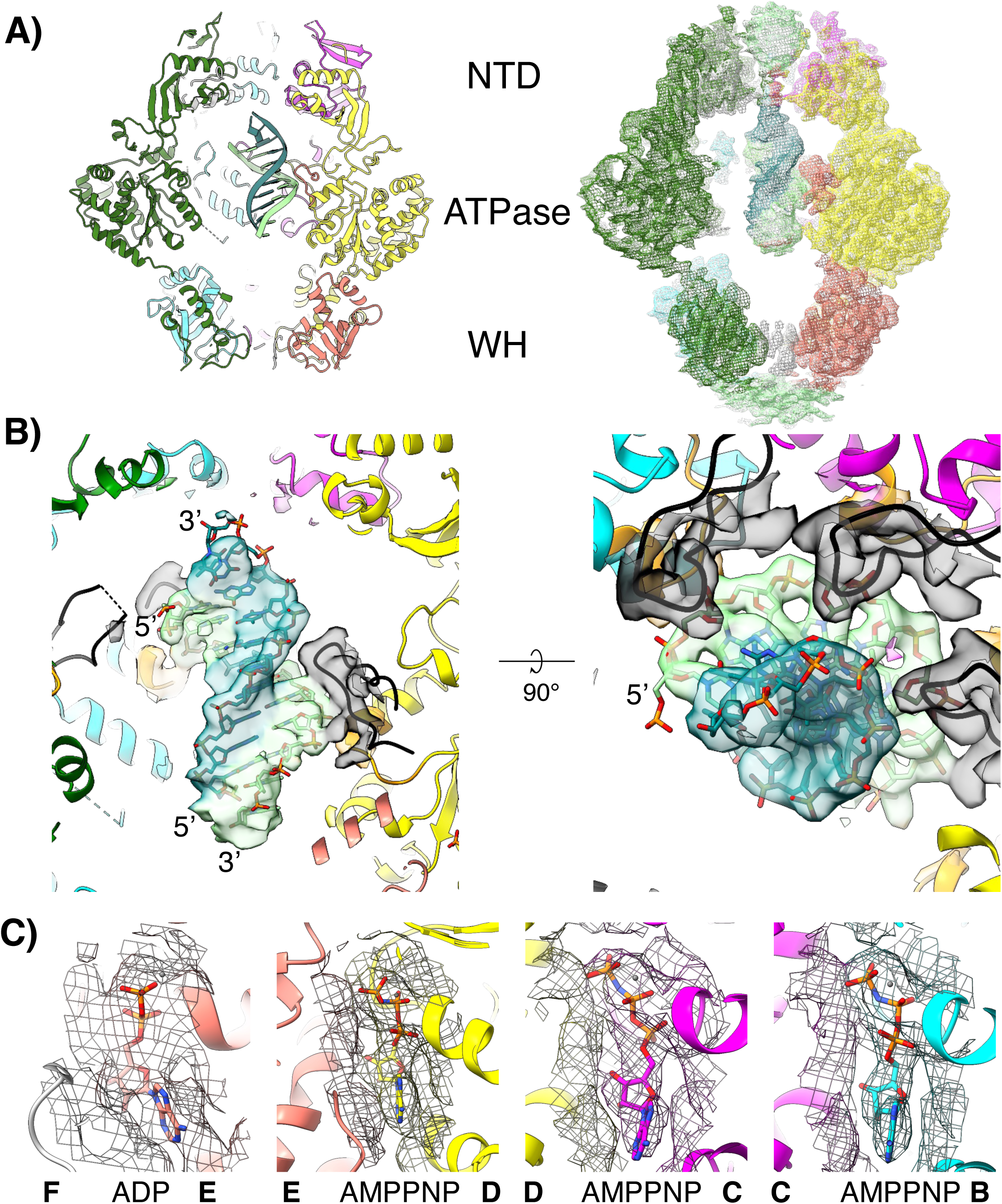
cryo-EM map of SaPIbov1 Rep. A. Cryo-EM structure of SaPIbov1 Rep (left) and map (right) shown in sliced side view, colored as previous figures. B. Close-up views of dsDNA density (contoured at 0.05) in sliced side (left) and top (right) views. C. Close-up views of ADP and AMPPNP densities, carved within 5Å of the nucleotides).

